# Glycogen metabolism jump-starts photosynthesis through the oxidative pentose phosphate pathway (OPPP) in cyanobacteria

**DOI:** 10.1101/657304

**Authors:** Shrameeta Shinde, Sonali P. Singapuri, Xiaohui Zhang, Isha Kalra, Xianhua Liu, Rachael M. Morgan-Kiss, Xin Wang

## Abstract

Cyanobacteria experience drastic changes in their carbon metabolism under daily light-dark cycles. In the light, the Calvin-Benson cycle fixes CO_2_ and divert excess carbon into glycogen storage. At night, glycogen is degraded to support cellular respiration. Dark-light transition represents a universal environmental stress for cyanobacteria and other photosynthetic lifeforms. Recent studies in the field revealed the essential genetic background necessary for the fitness of cyanobacteria during diurnal growth. However, the metabolic engagement behind the dark-light transition is not well understood. In this study, we discovered that glycogen metabolism can jump-start photosynthesis in the cyanobacterium *Synechococcus elongatus* PCC 7942 when photosynthesis reactions start upon light. Compared to the wild type, the glycogen mutant (Δ*glgC*) showed much lower photosystem II efficiency and slower photosystem I-mediated cyclic electron flow rate when photosynthesis starts. Proteomics analyses indicated that glycogen is degraded through the oxidative pentose phosphate pathway (OPPP) during dark-light transition. We confirmed that the OPPP is essential for the initiation of photosynthesis, and further showed that glycogen degradation through the OPPP is likely to contribute to the activation of key Calvin-Benson cycle enzymes by modulating NADPH levels during the transition period. This ingenious strategy helps jump-start photosynthesis in cyanobacteria following dark respiration, and stabilize the Calvin-Benson cycle under fluctuating environmental conditions. It has evolutionary advantages for the survival of photosynthetic organisms using the Calvin-Benson cycle for carbon fixation.

## Introduction

Photosynthesis supports life on earth by converting solar energy into chemical energy stored as organic carbon. The Calvin-Benson cycle, the major carbon fixation pathway in cyanobacteria, algae and plants, is employed to reduce CO2 into organic carbon molecules. Since its discovery in the 1950s, the enzymes of the Calvin-Benson cycle have been elucidated and consensus pathways established (Bassham et al., 1954). CO_2_ is reduced into the photosynthesis output glyceraldehyde 3-phosphate (G3P) in three stages, i.e. carbon fixation, carbon reduction, and carbon regeneration. The Calvin-Benson cycle is an elegant pathway that the starting substrate ribulose 1,5-bisphosphate (RuBP) is continuously supplied by partially recycling G3P back to RuBP through carbon rearrangement reactions. This carbon regeneration stage resembles the non-oxidative portion of the pentose phosphate pathway but with key differences. A key enzyme sedoheptulose-1,7-bisphosphatase (SBPase) was evolutionally selected to drive the carbon regeneration in the Calvin-Benson cycle rather than using the transaldolase found in the pentose phosphate pathway(Sharkey and Weise, 2016). The activity of SBPase often determines the photosynthetic capacity and carbon accumulation in downstream metabolic processes(Harrison et al., 1997; Raines et al., 1999; Harrison et al., 2001).

In cyanobacteria, a major portion of the photosynthesis-fixed carbon is used to synthesize the storage carbon glycogen. Two enzymes, ADP-glucose pyrophosphorylase (AGPase) and glycogen synthase (GS), catalyze the sequential conversion of glucose-1-phosphate (G1P) to 1,4-alpha-glucan (Preiss, 1984). Another enzyme encoded by 1,4-alpha-glucan-branching enzyme gene, *glgB*, can catalyze the formation of the 1,6-alpha branches of glycogen (Preiss, 1984). Glycogen synthesis is closely linked to photosynthesis carbon output. The photosynthate 3-phosphoglycerate (PGA) is an allosteric activator of the AGPase (Preiss, 1984; Gomez-Casati et al., 2003). When photosynthesis fixed carbon is in excess, a major carbon flux will be diverted to glycogen storage, which will serve as the carbon and electron sources for cellular respiration in the dark(Preiss, 1984; Suzuki et al., 2010). More recently, glycogen metabolism has also been recognized for additional roles played in cyanobacterial carbon metabolism. These studies revealed the involvement of glycogen metabolism as an energy buffering system to maintain homeostasis (Cano et al., 2018), and as the carbon source for rapid resuscitation from nitrogen chlorosis(Doello et al., 2018).

Glycogen metabolism and cellular respiration are critical for cell viability in the dark until light resumes(Lehmann and Wöber, 1976). Research in the past has identified the oxidative pentose phosphate (OPP) pathway, a.k.a. the glucose 6-phosphate (G6P) shunt, as the major route for glycogen degradation in the dark(Smith, 1983; Broedel and Wolf, 1990). On the other hand, the OPP pathway seems to become indispensable only after long period of dark (over 24 hours), while glycolysis might have been able to compensate for the need of reducing equivalent during short incubation in the dark(Scanlan et al., 1995). However, it is also important to know that OPP pathway is responsible for generating NADPH to serve as cofactors for certain ROS-detoxifying enzymes important for cyanobacterial survival (Welkie et al., 2018). In addition, cellular respiration is closely associated with photosynthesis. Many intermediates are shared between the carbon fixation and respiration processes. In cyanobacteria, many electron carriers in the thylakoid membrane are also shared between photosynthesis light reactions and cellular respiration(Mullineaux, 2014). When photosynthesis reactions start upon light, intermediates in the Calvin-Benson cycle might be limited after dark respiration, leading to stalled carbon fixation reactions. Dynamic metabolic regulation to ensure smooth transition from dark respiration to photosynthesis reactions is thus essential for the fitness of cyanobacteria during diurnal growth(Puszynska and O’Shea, 2017; Welkie et al., 2018). In this study, we discovered the active participation of glycogen metabolism during the dark-to-light transition period in cyanobacteria. We found that glycogen degradation through the OPP pathway helps activate and stabilize the Calvin-Benson cycle reactions when photosynthesis reactions start upon light. We propose that this ingenious strategy helps support photosynthesis during dark-to-light transition, and ensures the growth advantage of photosynthetic organisms in their natural environment.

## Results

### Proteomics analysis suggests a supportive role of glycogen metabolism for photosynthesis

Glycogen biosynthesis is a major carbon assimilatory pathway in the cyanobacterium *Synechococcus elongatus* PCC 7942. In this study, we generated a glycogen synthesis mutant by knocking out the AGPase gene (*glgC*) in the *S. elongatus* genome (Fig. 1A). When wild type and glycogen mutant cells were grown under moderate light levels (50 μmol photons m^−2^ s^−1^), the glycogen mutant exhibited a longer lag phase relative to wild type cells (Fig. 1B). This observation in the wild type cells suggests that glycogen metabolism might play a supportive role for the rapid photosynthesis start, i.e. a short lag phase to quickly start the carbon fixation process.

**Fig. 1.**
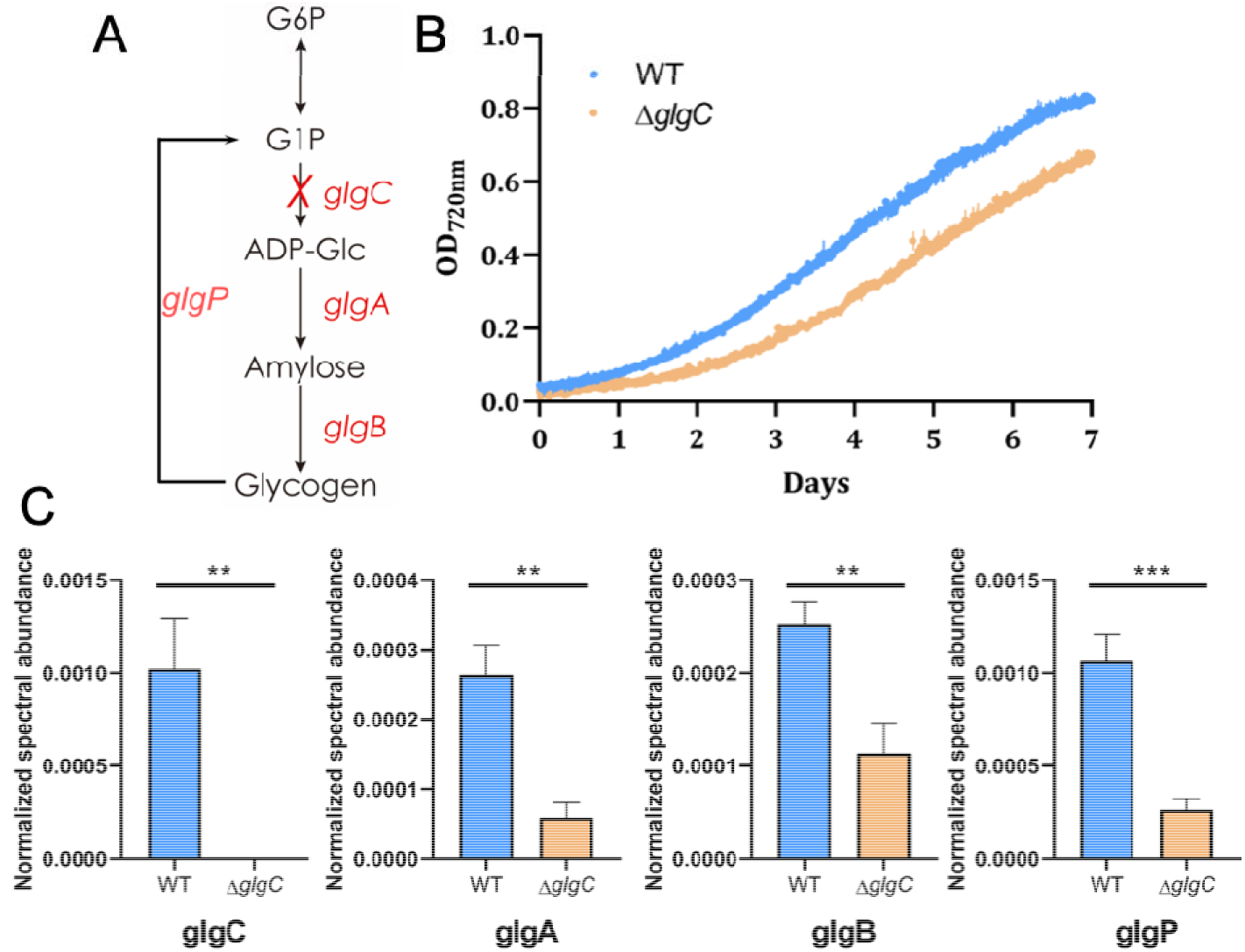
Impact of glycogen metabolism on *S. elongatus* growth. (A) Glycogen synthesis pathway in *S. elongatus*. The gene *glgC* was deleted from *S. elongatus* genome to create the glycogen mutant. (B) Growth curves of *S. elongatus* wild type and glycogen mutant cells (Δ*glgC*) measured by density optical at 720 nm. (C) Comparative proteomics analysis of *S. elongatus* wild type and glycogen mutant cells. Enzymes for both glycogen biosynthesis (AGPase (*glgC*), GS (*glgA*) and 1,4-alpha-glucan branching enzyme (*glgB*)) and degradation (alpha-1,4-glucan phosphorylase (glgP)) were highly expressed in wild type compared to glycogen mutant cells. Bar graph indicates enzyme expression levels based on the normalized spectral abundance factor (NSAF) (*p <0.05, **p<0.01, ***p<0.001).

To understand how glycogen biosynthesis supports photosynthesis, we conducted a comparative proteomics analysis on the wild type and glycogen mutant cells. The AGPase was not detected in the glycogen mutant cells, showing the success of constructing a clean glycogen mutant. All three enzymes related to glycogen synthesis, i.e. AGPase (glgC), glycogen synthase (*glgA*) and 1,4-alpha-glucan branching enzyme (*glgB*), were found to be expressed significantly higher in the wild type compared with those of glycogen mutant cells (Fig. 1C). Interestingly, enzymes related to glycogen degradation were also found to be more abundant in wild type cells. The alpha-1,4-glucan phosphorylase (*glgP*) that removes glucose from the glycogen chain was expressed over 4-fold higher in wild type cells (Fig. 1C and Table S1). However, if glycogen degradation were to continue through the glycolytic Embden-Meyerhof-Parnas (EMP) pathway, it would be a futile cycle to the carbon fixation pathway. We hypothesize that the glycogen hydrolysis might proceed through other alternative glycolytic pathways to support the rapid photosynthesis start in wild type cells. We began to test our hypothesis by determining the impact of abolished glycogen synthesis on photosynthetic function.

### The glycogen mutant has lower PSII efficiency when photosynthesis first starts

To test our theory that glycogen metabolism can support a higher photosynthesis efficiency when photosynthesis first starts, we measured the photosystem II (PSII) efficiency using the Phyto PAM II Walz fluorometer. When cyanobacterial cells were grown under continuous light, the maximum PSII efficiency (F_V_/F_M_) of wild type cells were relatively low in the first few hours of growth but gradually reached above 0.4, consistent with values reported for cyanobacteria(Campbell et al., 1998). The effective PSII efficiency (Φ_II_) exhibited a similar trend, with a measured efficiency ranging from 0.25 to 0.35 in the first 20 hours growing under continuous light conditions (Fig. 2A). Interestingly, the Φ_II_ of the glycogen mutant was significantly lower than that of wild type cells in the beginning of the growth phase, and only rose to a comparable level to the wild type cells at the later stage of growth (Fig. 2A). Thus, the differential levels of Φ_II_ between the wild type and glycogen mutant during the lag phase confirmed our hypothesis that glycogen biosynthesis has a supportive role for photosynthesis in *S. elongatus* when photosynthesis first starts.

**Fig. 2.**
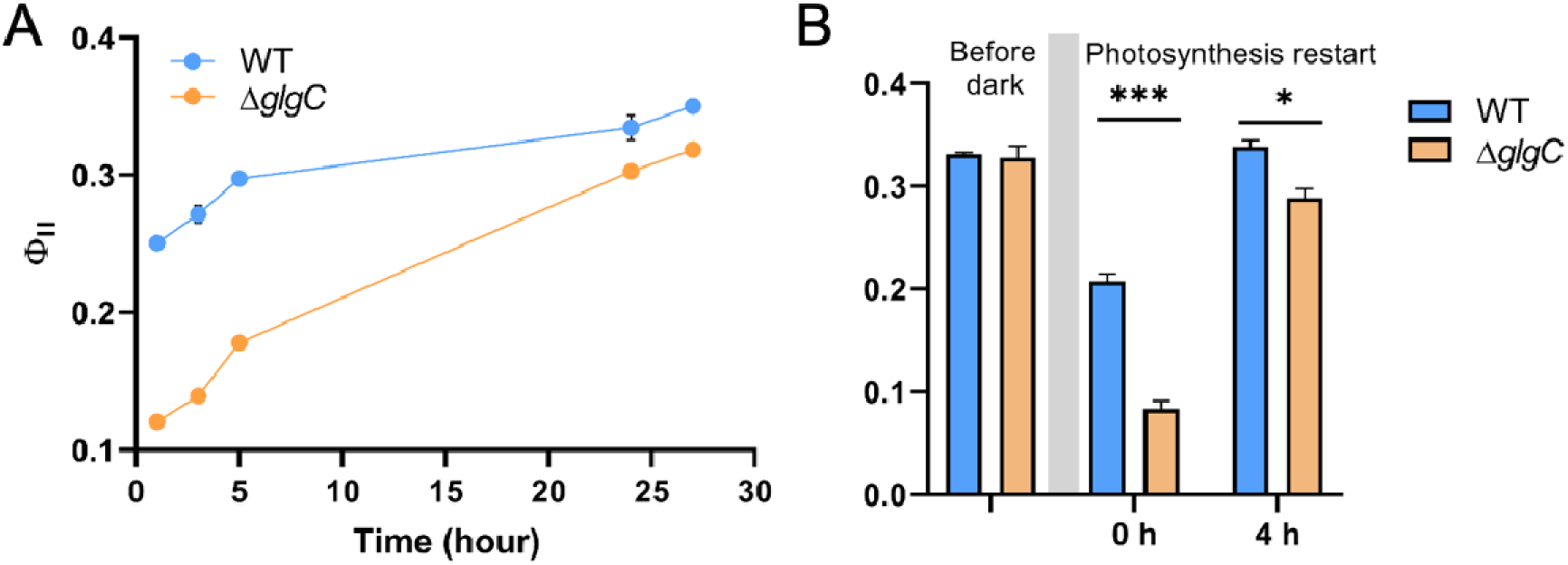
Photosystem II (PSII) efficiency of *S. elongatus* wild type and glycogen mutant cells. (A) The effective PSII efficiency (Φ_II_) measured under continuous light. Paired t test showed a p value of 0.0193 between wild type and glycogen mutant. (B) The effective PSII efficiency (Φ_II_) in dark incubated cells. Both Φ_II_ following the dark incubation (0 h) and after light was turned on for 4 hours were measured to monitor the effective PSII efficiency during photosynthesis restart. (*p<0.05, ***p<0.001)

To further validate the supportive role of glycogen metabolism in starting photosynthesis, we monitored Φ_II_ following prolonged dark incubation. Cellular respiration in the dark consumes the Calvin-Benson cycle intermediates, and gradually lowers or depletes the C_6_, C_5_ and C_3_ sugar intermediates pool(Iijima et al., 2015; Diamond et al., 2017). Without the support from glycogen metabolism, we thus should observe a lower photosynthetic efficiency in glycogen mutant cells when photosynthesis restarts after the light cycle resumes. To ensure the depletion of phosphate sugars, we incubated cells in dark for 30 hours. Following the dark incubation, the Φ_II_ was significantly higher in the wild type compared to the glycogen mutant cells during the first few hours of light (Fig. 2B), suggesting that PSII is transiently downregulated in the glycogen mutant. This result supports our hypothesis on the role of glycogen metabolism in sustaining a rapid start of photosynthesis.

### Photosystem I-mediated cyclic electron flow (CEF) is lower in the glycogen mutant

If the glycogen degradation goes through the alternative OPP pathway, there would be a higher ATP/NADPH ratio requirement during carbon fixation(Sharkey and Weise, 2016). Cyclic electron flow (CEF) around photosystem I (PSI) contributes largely to additional ATP requirements in cyanobacterial carbon metabolism (Mullineaux, 2014). We thus measured the CEF rate through P700 photooxidation under far-red (FR) light(Klughammer and Schreiber, 1994; Cook et al., 2019). Oxidized PSI (P700^+^) absorbs strongly at wavelengths between 810-820 nm (A_820_), while reduced P700 has minimal absorbance at this range. Both wild type and glycogen mutant cells exhibited a rapid increase in A820 after the FR light was turned on, indicating the oxidation of PSI (P700 to P700^+^). Once a stable A820 was attained, the FR was turned off, and the rate of re-reduction of P700^+^ to P700 (*t_½_^red^*) was calculated as an estimate of P700^+^ re-reduction from alternative electron donors, of which CEF is the major pathway(Xu et al., 1994). We monitored CEF rates in wild type and glycogen mutant cultures under both dark and light conditions. P700 kinetics was monitored in log-phase *S. elongatus* wild type and glycogen mutant cells both at the end of dark incubation and after light was turned on for 2 hours and 24 hours. The ratio of ΔA_820_/A_820_ was comparable for both strains under all conditions (Fig. 3), indicating that, unlike PSII, the amount of photooxidizable P700 was not affected by the mutation in the glycogen synthesis pathway. After dark incubation for 16 hours, both the wild type and glycogen mutant cells exhibited relatively slow *t_½_*^red^, suggesting that CEF is low in the cyanobacterial cultures following dark incubation (Fig. 3). However, the wild type cells exhibited more than a 2-fold faster *t_½_*^red^ after 2 hours of incubation in the light. After 24 hours into the light, the *t_½_*^red^ of both the wild type and glycogen mutant recovered to comparable levels (Fig. 3).

**Fig. 3.**
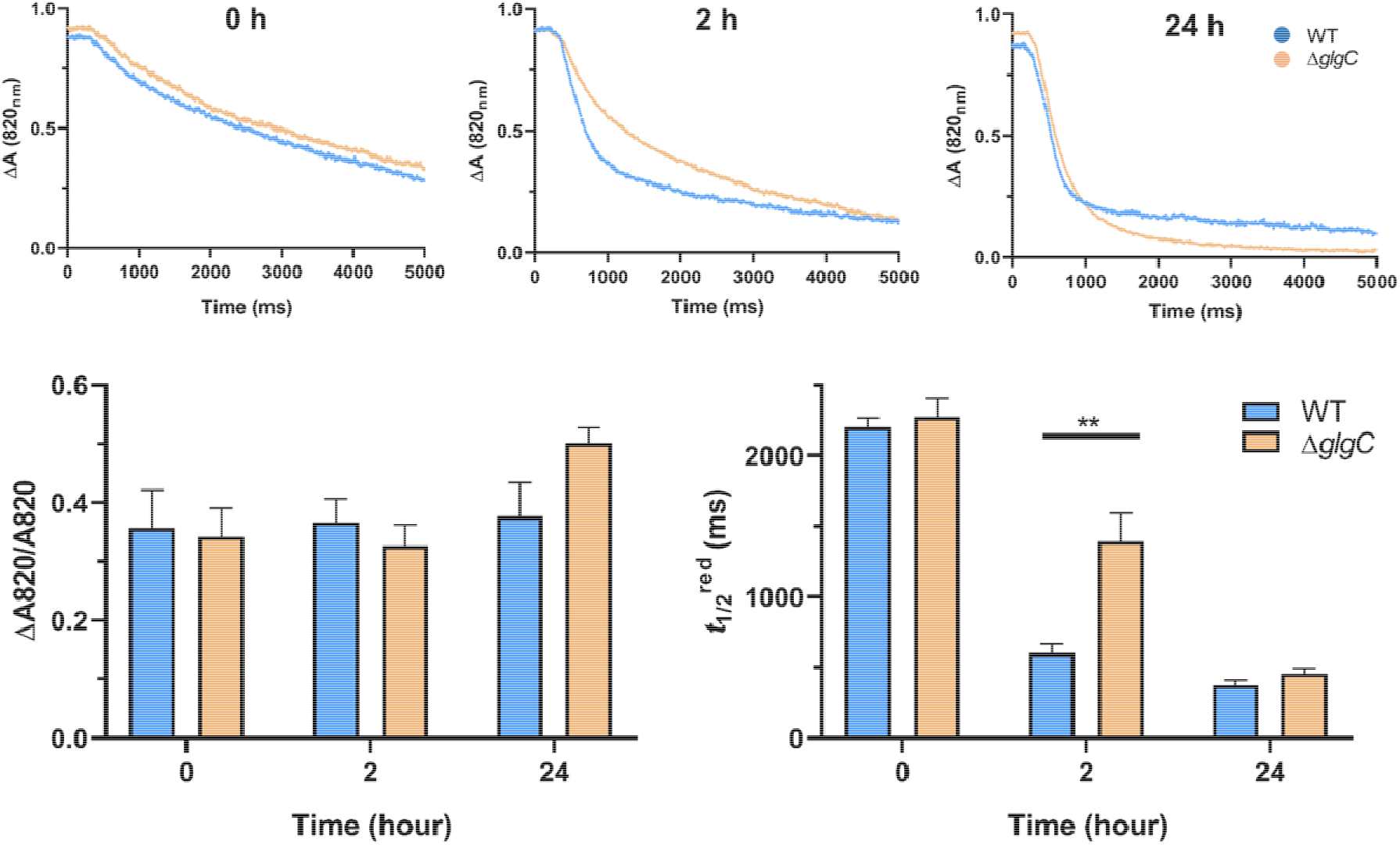
P700^+^ reduction in *S. elongatus* wild type and glycogen mutant cells following dark incubation. Cultures were incubated in the dark for 16 hours. The P700^+^ reduction was measured right after dark adaption (0 h), 2 hours into the light (2 h), and 24 hours into the light (24 h). The data was plotted from the means of 9 replicates (3 biological × 3 technical replicates). Bar graphs indicate the oxidizable P700 pool (ΔA820/A820) and the calculated P700^+^ re-reduction rate (*t½*^red^) from the slope of the reduction curve. (**p<0.01).

### Glycogen metabolism jump-starts photosynthesis through the oxidative pentose phosphate pathway

To determine how glycogen metabolism can help jump-start photosynthesis, we conducted another comparative proteomics study on dark-incubated cyanobacterial cells. Log-phase wild type and glycogen mutant cells were dark incubated for 16 hours before the light was turned on. After two hours into the light, cells were collected for proteomics analysis. Similar to log-phase cells collected under continuous light, the enzymes for glycogen metabolism, i.e. AGPase (*glgC*, glycogen synthase (*glgA*), 1,4-alpha-glucan branching enzyme (*glgB*), and alpha-1,4-glucan phosphorylase (glgP), were found in significantly higher abundances in wild type cells compared to those of glycogen mutant cells (Fig. 4A). Interestingly, the enzyme 6-phosphogluconate dehydrogenase (*gnd*), a key enzyme of the OPP pathway, was also expressed significantly higher in wild type cells (Fig. 4A). The proteomics analysis indicates that the glycogen degradation might proceed through the alternative OPP pathway.

**Fig. 4.**
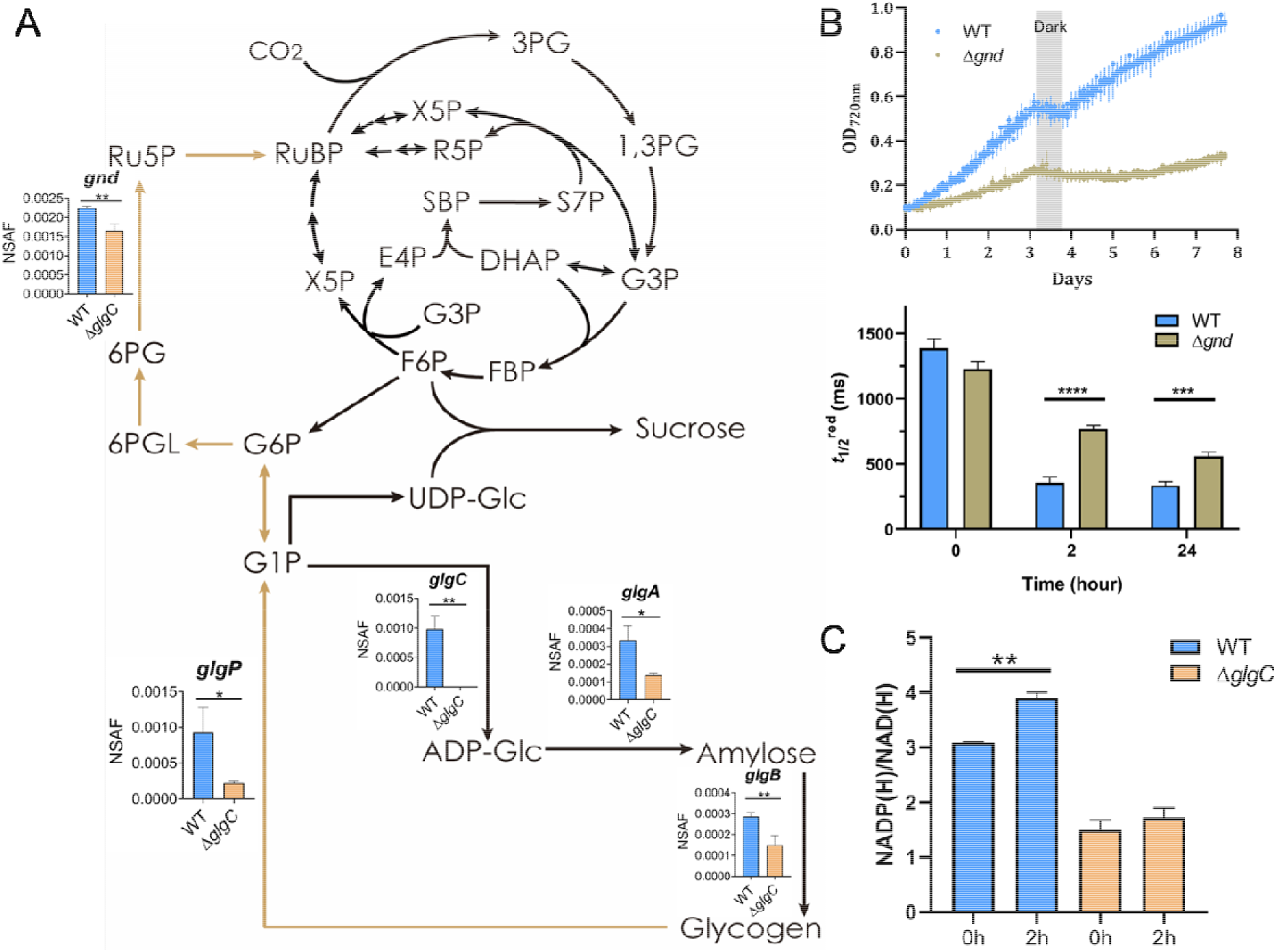
Glycogen metabolism jump-starts photosynthesis through the OPPP. (A) Yellow arrows indicate the proposed mechanism in which glycogen degrades through the OPPP to recharge photosynthesis during the start of photosynthesis. Bar graphs indicate the protein levels from proteomics analysis. (B) Growth profile and P700^+^ re-reduction analysis of wild type and the OPP pathway mutant Δ*gnd*. Bar graph indicates the calculated P700^+^ re-reduction rate (*t_½_*^red^). (C) Detection of NADP(H)/NAD(H) ratio in both wild type and glycogen mutant Δ*glgC* cells following dark incubation and after 2 hours into the light. (*p <0.05, **p<0.01, ***p<0.001). (**Relevant acronyms**: **3PG**: 3-phosphoglycerate; **G3P**: Glyceraldehyde 3-phosphate; **Ru5P**: Ribulose 5-phosphate; **RuBP**: Ribulose 1,5-bisphosphate; **G6P**: Glucose 6-phosphate; **6PGL**: 6-phosphogluconolactone; **G1P**: Glucose 1-phosphate; **ADP-Glc**: ADP-glucose.)

To validate the involvement of the OPP pathway in stimulating a rapid photosynthesis start, we generated an OPP pathway mutant strain (Δ*gnd*) by deleting the 6-phosphogluconate dehydrogenase gene (*gnd*) in the *S. elongatus* genome. The selection of *gnd* over the G6P dehydrogenase gene (*zwf*) for the deletion is to minimize the potential impact on other metabolic process such as the Entner-Doudoroff pathway in need of the G6P dehydrogenase to proceed (Chen et al., 2016). Similar to the glycogen mutant cells, the Δ*gnd* strain showed impaired growth compared to wild type cells when photosynthesis starts (Fig. 4B). We further measured the P700^+^ re-reduction rate of the Δ*gnd* cells. The wild type and Δ*gnd* cells were grown to log phase under moderate light levels (50 μmol photons m^−2^ s^−1^) before the light was turn off for 16 hours. Following transition to light, the P700^+^ re-reduction rates of both wild type and Δ*gnd* cells were slow after dark incubation (Fig. 4B). However, the *t_½_*^red^ of wild type cells quickly recovered after 2 hours of incubation in the light, whereas the *t_½_*^red^ of the Δ*gnd* cells was significantly slower (Fig. 4B). Even after 24 hours in light, the *t_½_*^red^ of the Δ*gnd* cells was still relatively slow compared to wild type cells, indicating long-term downregulation of photosynthesis (Fig. 4B). This observation confirms our hypothesis that the OPP pathway is crucial for the rapid photosynthesis start after dark incubation, and the healthy performance of photosynthesis in the light.

### The OPP pathway modulates the NADP(H)/NAD(H) levels to activate the Calvin-Benson cycle

Upon restarting of photosynthesis reactions following dark incubation, the intermediates pool needed to regenerate RuBP in the Calvin-Benson cycle may be limited or depleted(Iijima et al., 2015), leading to inferior photosynthesis performance and downregulation of the photosynthetic electron transport chain. Thus, continuous supply of RuBP during the dark-light transition is likely critical for maintaining homeostatic balance between energy production by the light reactions and energy consumption by the Calvin-Benson cycle. The obvious outcome for the glycogen degradation through the OPPP is to replenish C5 carbon pools. Ru5P derived from the OPP pathway can then be phosphorylated to RuBP by the phophoribulokinase (PRK), replenishing the starting substrate for RuBisCO to continuously fix CO_2_ in the Calvin-Benson cycle. However, PRK is inactive in the dark and requires redox regulation to become active(Tamoi et al., 2005). To validate whether the OPP pathway help activate photosynthesis, we measured the levels of reducing equivalents in wild type and glycogen mutant cells following dark incubation and after light was turned on for two hours. When photosynthesis reactions restart after dark incubation, the NADP(H)/NAD(H) ratio was significantly higher in the wild type than the glycogen mutant cells (Fig. 4C), supporting our hypothesis that glycogen metabolism through the OPP pathway can help activate the Calvin-Benson cycle for a rapid start of photosynthesis.

## Discussion

Photosynthetic organisms experience drastic changes in their carbon metabolism during diurnal growth. Recent research has highlighted the contribution of glycogen metabolism and the OPP pathway to the fitness of cyanobacteria during diurnal growth (Diamond et al., 2017; Puszynska and O’Shea, 2017; Welkie et al., 2018). However, knowledge on metabolic engagement during the initiation of photosynthesis is not well understood. This study emphasizes the active role played by the glycogen metabolism and the OPP pathway when photosynthesis starts after dark respiration. Our results indicate that this supportive role is likely essential for the survival of cyanobacteria in their natural environment.

Cellular respiration from glycogen degradation helps maintain the cell integrity in dark. When light resumes, many enzymes for cellular respiration are inactivated, shifting cell metabolism to carbon fixation(Udvardy et al., 1984; Gleason, 1996; Sharkey and Weise, 2016). However, several recent studies have shown that cellular respiration might play more important roles than just maintaining cell viability for dark survival. A glycogen phosphorylase (Δ*glgP*) mutant of *Synechocystis* sp. PCC 6803 retards dark respiration of glycogen, and had a much lower photosynthetic oxygen evolution rate(Shimakawa et al., 2014). Another study conducted in the glycogen mutant Δ*glgC* of *Synechocystis* sp. PCC 6803 also exhibited a delayed activation of the Calvin-Benson cycle, and diminished photochemical efficiency when cultured under high carbon conditions (Holland et al., 2016). These studies indicate a tight link between respiration and the photosynthesis initiation process. In this study, we showed that glycogen metabolism can recharge photosynthesis through the OPP pathway in the unicellular cyanobacterium *S. elongatus* when photosynthesis reactions start upon light (Fig. 4). When light resumes after dark incubation, RuBP concentration is low due to the suppressed PRK activity in the dark(Tamoi et al., 2005). This alternative carbon flow is thus essential to stabilize photosynthesis when triose phosphate (TP) levels are insufficient to replenish RuBP through the Calvin-Benson cycle during the start of photosynthesis. The importance of the OPP pathway is further manifested by the lack of efficiency to synthesize TPs through glycogen degradation via phosphofructokinase (PFK) and phosphoglucoisomerase (PGI), both of which require high level of substrate (Knowles and Plaxton, 2003; Preiser et al., 2018). Glycogen metabolism thus serves as a perfect buffering system to supply G6P to the shunt by sensing the photosynthesis output PGA accumulation under the condition of stalled Calvin-Benson cycle reactions (Gomez-Casati et al., 2003). In addition, our proteomics results also showed that the transaldolase (*tal*), a non-essential enzyme in the reductive phase of the pentose phosphate pathway, was expressed significantly higher in the wild type compared to the glycogen mutant cells when photosynthesis reactions first start (Table S1). This result suggests an underappreciated role for transaldolase during the operation of the OPPP. Interestingly, a recent genome-wide fitness study in *S. elongatus* showed similar result that the *tal* mutant was sensitive to light-dark cycles(Welkie et al., 2018).

Our findings echoes with the proposed mechanism by Sharkey et al. for the involvement of the OPP pathway to stabilize photosynthesis in plants (Sharkey and Weise, 2016). Sharkey suggested that the OPP pathway (a.k.a. G6P shunt) could account for 10-20% of Rubisco activity, and the enzyme SBPase plays an important role in controlling the carbon flow between the Calvin-Benson cycle and the shunt(Sharkey and Weise, 2016). Their recent evidence showed that G6P shunt was activated in a photorespiration mutant of *Arabidopsis thaliana* in which the activity of triose phosphate isomerase (TPI) was inhibited(Li et al., 2018). TPI is responsible for interconverting G3P and dihydroxyacetone phosphate (DHAP), a key reaction for RuBP regeneration in the Calvin-Benson cycle. During photosynthesis, it is critical that RuBP can be continuously regenerated for CO2 fixation, which limits the carbon regeneration reactions to a small margin of error in order to maintain the stable operation of photosynthesis. Perturbation to the carbon regeneration stage of the Calvin-Benson cycle such as the TPI inhibition would lead to a temporarily impaired Calvin-Benson cycle. An alternative mechanism such as the G6P shunt is thus essential to stabilize photosynthesis.

The operation of the OPPP during the start of photosynthesis was revealed by the energy generation from photosynthetic light reactions. Compared to the wild type, the PSII efficiency in the glycogen mutant was significantly lower when photosynthesis starts (Fig. 3), suggesting that PSII is likely downregulated through a feedback mechanism in the glycogen mutant. We also observed different P700^+^ re-reduction rates in the wild type compared to both mutant strains (Δ*glgC* and Δ*gnd*). Both wild type and glycogen mutant cells had comparably slow P700^+^ re-reduction rate after dark incubation; however, wild type exhibited rapid recovery whithin 2 hours in light, while it took 24 hours for the glycogen mutant to reach a comparable level to the wild type cells (Fig. 3). When comparing between wild type and the Δ*gnd* mutant, the P700^+^ re-reduction rate showed similar trends, i.e. a much faster re-reduction rate in wild type cells after a short incubation in the light (Fig. 4B). This strongly suggests a higher ATP/NADPH ratio requirement when photosynthesis first starts and is directly linked to glycogen metabolism and the OPP pathway. The action of OPP pathway along the Calvin-Benson cycle exacerbates the deficit of ATP/NADPH ratio needed for carbon fixation(Sharkey and Weise, 2016). It is well accepted that the additional ATP requirement in photosynthetic carbon metabolism can be fulfilled through the cyclic electron flow(Kramer and Evans, 2011). The higher CEF rate observed in the wild type cells fits well with the additional ATP requirements for operating the OPPP during the start of photosynthesis. Compared to the glycogen mutant, the Δ*gnd* mutant did not recover to a comparable level of P700^+^ re-reduction rate even after 24 hours into the light, indicating that the OPP pathway is essential for the initiation of photosynthesis and the fitness of cells (Fig. 4B). When the Calvin-Benson cycle reactions are fully operational (i.e. hours in the light), the glycogen metabolism and the OPPP might play smaller roles, as revealed by the similar P700^+^ re-reduction rates in light-adapted wild type and glycogen mutant cells (Fig. 3).

Glycogen degradation through the OPP pathway could replenish RuBP to ensure the continuous operation of carbon fixation during the start of photosynthesis. However, it is important to know that the OPP pathway does not lead to any net carbon fixation. The CO2 fixed in the Calvin-Benson cycle is quickly lost when G6P is oxidized to Ru5P in the shunt. It is thus intriguing to understand the role the OPP pathway in supporting photosynthesis. We propose that the OPP pathway could generate additional NADPH required for the redox regulation of the Calvin-Benson cycle during the start of photosynthesis. Previous research showed that the activities of two key enzymes in the Calvin-Benson cycle, PRK and G3P dehydrogenase (GADPH), are suppressed in dark through the formation of a CP12/PRK/GAPDH protein complex(Tamoi et al., 2005). The reversible dissociation of the complex was controlled by the NADP(H)/NAD(H) ratio in *S. elongatus*(Tamoi et al., 2005). The operation of the OPP pathway during the start of the photosynthesis thus could contribute largely to the increase of NADPH levels when light resumes, releasing PRK and GAPDH from the complex to activate the Calvin-Benson cycle. The measurement of reducing equivalents both at the end of dark incubation and after light was turned on for two hours supports this hypothesis. Compared to the glycogen mutant Δ*glgC* cells, the increase of NADP(H)/NAD(H) ratio in the wild type was significantly larger after the light was turned on, i.e. when photosynthesis restarts (Fig. 4C).

Lastly, it is also important to understand the activation mechanism of the OPP pathway. The expression of many genes in the glycogen metabolism and the OPP pathway are controlled by the circadian clock output protein RpaA(Markson et al., 2013; Welkie et al., 2018). Our proteomics analysis confirmed that many enzymes in these two metabolic processes were expressed much higher in wild type cells in the beginning of photosynthesis (Fig. 4A and Table S1). Previous research also reported that the first enzyme G6P dehydrogenase (G6PDH) in the OPP pathway is redox regulated and is inactive in its reduced form(Udvardy et al., 1984; Gleason, 1996). However, G6P could also stabilize and activate G6P dehydrogenase to reverse the effect of reduction inhibition in cyanobacteria(Cossar et al., 1984). It is thus possible that glycogen degradation would lead to increased G6P levels and activate the G6P dehydrogenase. Future studies toward understanding these mechanisms could help fully appreciate the roles played by the OPP pathway in supporting photosynthesis.

In conclusion, we found that glycogen metabolism in cyanobacteria could help activate photosynthesis through the OPP pathway during the start of photosynthesis reactions. The OPP pathway is generally considered for its role to generate reducing equivalent for dark respiration. However, evolution empowers cells with the ability to leverage existing metabolic processes to its advantage in order to adapt to the constantly changing environment. We have witnessed an interesting strategy implemented by cyanobacteria to ensure their growth advantage in their natural environment.

## Materials and Methods

### Growth conditions

Seed cultures of *S. elongatus* wild type and mutant strains were grown in BG11 media supplemented with 20 mM sodium bicarbonate and 10 mM N-[Tris(hydroxymethyl)methyl]-2-aminoethanesulfonic acid (TES, pH 8.2) at 30 °C under 30 μmol photons m^−2^ s^−1^ illumination. The growth media of the mutant strains were also supplemented with 5 mg/L kanamycin. Cultures for measurements were grown at 30 °C under 50 μmol photons m^−2^ s^−1^ illumination in the Multi-Cultivator MC-1000-OD-MULTI (Photon Systems Instruments, Czech Republic). Cyanobacterial growth was monitored using the built-in spectrophotometer of Multi-Cultivator by measuring optical densities at the wavelength of 720 nm.

### Plasmid and strain construction

The gene fragments for plasmid construction were amplified using primers designed through the NEBuilder Assembly tool (http://nebuilder.neb.com/). The *glgC* and *gnd* knock out plasmids were constructed using HiFi DNA Assembly master mix (NEB, Ipswich, MA). *S. elongatus* wild type cells were transformed with knockout plasmids to generate Δ*glgC* and Δ*gnd* mutant strains. The strains and plasmids along with primers used in this study are listed in the supplemental information in Table S2 and S3, and File S1 and S2.

### Proteomics Analysis

Log-phase *S. elongatus* wild type and glycogen mutant cells were used for proteomics analysis following our previously described method (Wang et al., 2016). For dark-incubated samples, log-phase cells were taken after light was turned on for two hours following dark incubation. Briefly, biological triplicates of approximate 10 OD of cells were used for protein extraction. Cell pellets were resuspended in 1 mL of Tris buffer (50 mM Tris-HCl, 10 mM CaCl_2_, 0.1 % n-Dodecyl β-D-maltoside, pH 7.6), followed by cell lysis using a homogenizer with bead-beating cycles of 10 s ON and 2 min OFF on ice for a total of 6 cycles. Protein extract in the supernatant was collected by centrifugation at 13000g for 30 min at 4°C, followed by protein concentration measurement using the Bradford assay (Thermo Scientific).

For each sample, 100 μg of total protein was digested by Trypsin Gold (Promega, Madison, WI) with 1:100 w/w ratio at 37 °C for 18 hours. The digested peptides were cleaned up using a Sep-Pak C18 column (Waters Corporation, Milford, MA), followed by peptide fractionation using the Pierce High pH Reverse-Phase Peptide Fractionation Kit (Thermo Scientific, Rockford, IL). Eight fractions of peptides from each sample were subjected to liquid chromatography-tandem mass spectrometry (LC-MS/MS) analysis in a Thermo LTQ Orbitrap XL mass spectrometer. The mass spec analysis was operated under the data-dependent mode scanning the mass range of 350-1800 *m/z* at the resolution of 30,000. The 12 most abundant peaks were subjected to MS/MS analysis by the collision induced dissociation fragmentation. The raw data collected from MS/MS analysis was searched and analyzed using the pipeline programs integrated in the PatternLab for Proteomics (version 4.1.0.17)(Carvalho et al., 2016). The normalized spectral abundance factor (NSAF) was used to compare protein abundance in different groups(Zybailov et al., 2006). The mass spectrometry proteomics data have been deposited to both the MassIVE repository with the dataset identifier MSV000083575, and the ProteomeXchange Consortium(Vizcaíno et al., 2015) with the dataset identifier PXD013099.

### Photosystem II (PSII) analysis

Cyanobacterial cells grown both under continuous light (50 μmol photons m^−2^ s^−1^) and dark incubated cells were subjected to PSII measurement using the Phyto-PAM Phytoplankton Analyzer (Walz, Germany). When measuring PSII efficiency, the dark adaption that normally oxidizes the PSII center in green algae does not work in *S. elongatus*. Instead, cells would transit to state II after dark adaptation(Campbell et al., 1998). To get a proper measurement of PSII efficiency, cells were illuminated at moderate actinic light (25 μmol/m^2^/s) for 5 minutes to lock the photosystems in state I and to oxidize PSII reaction centers. Standard induction curves by exposing cells to saturation pulses were recorded for the measurement of the maximum PSII efficiency (F_V_/F_M_) and the effective PSII efficiency (Φ_II_).

### Photosystem I (PSI) analysis

Log-phase *S. elongatus* wild type and mutant cells were incubated in dark for 16 hours before the PSI analysis. Five mL of the culture was vacuum-filtered through Whatmann GF/C 25 mm filter discs (Cat# 1822-025). Far red light induced P700 oxidation-reduction kinetics measurements were performed using a dual-wavelength pulse amplitude-modulated fluorescence monitoring system (Dual-PAM-100, Heinz Walz, Effeltrich, Germany) with a leaf attachment. The proportion of photooxidizable P700 (Δ*A820/A820*) was determined as the change in absorbance at 820 nm after turning on the far-red light *λ*_max_□=□715 nm, 10 Wm^−2^, Scott filter RG 715). The half-time for the re-reduction of P700^+^ to P700 (*t_½_*^red^) was calculated after the far-red light was turned off and used as an estimate of alternative electron flow around PSI.

### Measurement of NADP(H) and NAD(H) levels

Log-phase *S. elongatus* wild type and Δ*glgC* cells were incubated in the dark for 16 hours before the light was turned on. 50 μL of cells were collected in triplicates both at the end of dark incubation and after the light was on for 2 hours. The levels of pyridine nucleotides NAD(P)^+^ and NAD(P)H were determined by the NAD/NADH-Glo™ and NADP/NADPH-Glo™ assays (Cat# G9071, Promega, Madison, WI) following the manufacturer s instructions.

## Acknowledgements

This work was supported by Miami University startup fund to XW. XW and RMK are also supported by Department of Energy-BES Photosynthetic Systems program. We thank Drs. Spencer Diamond, James Golden, and Susan Golden for the original plasmid used to generate the glycogen mutant strain in the preliminary study. We would like to thank Dr. Theresa Ramelot and Kundi Yang of the chemistry department at Miami University for their help in the preliminary metabolite analysis. We thank Dr. Jianping Yu (NREL) for the careful reading of the manuscript.

## Author contributions

XW designed and performed the experiment, analyzed data, and wrote the manuscript. SS, SPS, XZ and IK performed the genetics, proteomics, and photochemistry experiments. RMK helped analyze the photochemistry data. SS, XL and RMK edited the manuscript.

