## Supplemental Information for "Glycogen metabolism jump-starts photosynthesis through the oxidative pentose phosphate pathway (OPPP) in cyanobacteria"

Supplementary Table 1-3
Supplementary Files 1-2

Table S1. Comparative proteomics analysis of *S. elongatus* wide type and Δ*glgC* cells. Results from cells grown under both continuous light and dark-adapted conditions were summarized in the table. The listed proteins are the candidate proteins showing statistical differences in their abundances by comparing between wild type (WT) and glycogen mutant cells.

**Group 1 (Continuous light):**
Identifications satisfied both the automatic fold and statistical criteria in the Patternlab analysis.

| UniProt ID | Fold Change | p Value | Δ*glgC* | WT | Description |
|---|---|---|---|---|---|
| Q31RP3 | 4.072368 | 0.000437 | 0.000262 | 0.001066 | Alpha-1,4 glucan phosphorylase OS=*Synechococcus elongatus* (strain PCC 7942) GN=Synpcc7942_0244 PE=3 SV=1 |
| Q31RP2 | 2.866004 | 0.029407 | 0.000236 | 0.000677 | Glyceraldehyde-3-phosphate dehydrogenase OS=*Synechococcus elongatus* (strain PCC 7942) GN=Synpcc7942_0245 PE=3 SV=1; Additional IDs concatenated into MaxParsimony group: Reverse_Q31KI0_PURA_SYNE7 |
| Q31KD7 | 1.930478 | 0.002943 | 0.000353 | 0.000681 | Tfp pilus assembly protein PilN-like OS=*Synechococcus elongatus* (strain PCC 7942) GN=Synpcc7942_2452 PE=4 SV=1 |
| P0A3D2 | 1.844583 | 0.041368 | 0.00803 | 0.014812 | Ferredoxin-1 OS=*Synechococcus elongatus* (strain PCC 7942) GN=petF PE=3 SV=2 |
| Q31LW3 | 1.839106 | 0.018559 | 0.000375 | 0.000689 | Uncharacterized protein OS=*Synechococcus elongatus* (strain PCC 7942) GN=Synpcc7942_1926 PE=4 SV=1 |
| Q31LL0 | 1.732225 | 0.037129 | 0.000655 | 0.001134 | Glucose-6-phosphate isomerase OS=*Synechococcus elongatus* (strain PCC 7942) GN=pgi PE=3 SV=1 |
| Q31KD8 | 1.640014 | 0.01924 | 0.000367 | 0.000603 | Putative type IV pilus assembly protein PilO OS=*Synechococcus elongatus* (strain PCC 7942) GN=Synpcc7942_2451 PE=4 SV=1 |
| Q31QA2 | 1.569555 | 0.015596 | 0.00044 | 0.000691 | Uncharacterized protein OS=*Synechococcus elongatus* (strain PCC 7942) GN=Synpcc7942_0735 PE=4 SV=1 |
| Q31P93 | 1.425252 | 0.00657 | 0.000562 | 0.000801 | Phosphomethylpyrimidine synthase OS=*Synechococcus elongatus* (strain PCC 7942) GN=thiC PE=3 SV=1 |
| Q31Q47 | -1.72735 | 0.003124 | 0.004981 | 0.002884 | RNA-binding region RNP-1 OS=*Synechococcus elongatus* (strain PCC 7942) GN=Synpcc7942_0790 PE=4 SV=1 |
| Q31S60 | -1.83673 | 0.000439 | 0.000605 | 0.000329 | Transcriptional regulator, XRE family OS=*Synechococcus elongatus* (strain PCC 7942) GN=Synpcc7942_0077 PE=4 SV=1 |

| | | | | | |
|---|---|---|---|---|---|
| Q31PN2 | -2.20224 | 0.001754 | 0.000685 | 0.000311 | Cob(I)yrinic acid a,c-diamide adenosyltransferase OS=*Synechococcus elongatus* (strain PCC 7942) GN=Synpcc7942_0957 PE=4 SV=1 |

**Group 2 (Continuous light):**
These identifications were filtered out by the L-stringency in the Patternlab analysis.

| UniProt ID | Fold Change | p Value | Δ*glgC* | WT | Description |
|---|---|---|---|---|---|
| Q31M00 | 4.736188 | 0.006 | 6.76E-05 | 0.00032 | Uncharacterized protein OS=*Synechococcus elongatus* (strain PCC 7942) GN=Synpcc7942_1889 PE=3 SV=1 |
| Q935Y7 | 4.499441 | 0.000879 | 5.87E-05 | 0.000264 | Glycogen synthase OS=*Synechococcus elongatus* (strain PCC 7942) GN=glgA PE=3 SV=3 |
| Q31L64 | 3.517046 | 0.019264 | 7.35E-05 | 0.000259 | Iron deficiency-induced protein A OS=*Synechococcus elongatus* (strain PCC 7942) GN=idiA PE=1 SV=2 |
| Q31MW7 | 3.132478 | 0.004709 | 9.80E-05 | 0.000307 | Dehydrogenase subunit-like protein OS=*Synechococcus elongatus* (strain PCC 7942) GN=Synpcc7942_1572 PE=4 SV=1 |
| Q31N05 | 2.771257 | 0.049696 | 6.97E-05 | 0.000193 | Uncharacterized protein OS=*Synechococcus elongatus* (strain PCC 7942) GN=Synpcc7942_1534 PE=4 SV=1 |
| Q31NW5 | 2.465642 | 0.002982 | 8.96E-05 | 0.000221 | ABC-transporter membrane fusion protein OS=*Synechococcus elongatus* (strain PCC 7942) GN=Synpcc7942_1224 PE=4 SV=1 |
| Q31M40 | 2.431201 | 0.023822 | 0.000144 | 0.000349 | RNA polymerase sigma factor OS=*Synechococcus elongatus* (strain PCC 7942) GN=Synpcc7942_1849 PE=3 SV=1; Additional IDs concatenated into MaxParsimony group: Q31QG5_SIGA3_SYNE7, Q31QR8_SIGA4_SYNE7 |
| P16954 | 2.226871 | 0.002183 | 0.000113 | 0.000253 | 1,4-alpha-glucan branching enzyme GlgB OS=*Synechococcus elongatus* (strain PCC 7942) GN=glgB PE=1 SV=2 |
| Q31KP0 | 2.218107 | 0.03169 | 0.00024 | 0.000533 | Twitching motility protein OS=*Synechococcus elongatus* (strain PCC 7942) GN=Synpcc7942_2349 PE=4 SV=1 |
| Q31PQ0 | 2.105309 | 0.020686 | 0.000159 | 0.000335 | Adenylyl-sulfate kinase OS=*Synechococcus elongatus* (strain PCC 7942) GN=cysC PE=3 SV=1 |
| Q31KE8 | 2.08357 | 0.009504 | 0.000208 | 0.000433 | Phosphate import ATP-binding protein PstB OS=*Synechococcus elongatus* (strain PCC 7942) GN=pstB PE=3 SV=1 |

| | | | | | |
|---|---|---|---|---|---|
| Q31Q75 | 1.954468 | 0.002478 | 0.000281 | 0.000548 | Uncharacterized protein OS=*Synechococcus elongatus* (strain PCC 7942) GN=Synpcc7942_0762 PE=4 SV=1 |
| Q31S25 | 1.944628 | 0.027214 | 7.41E-05 | 0.000144 | Deoxyribodipyrimidine photo-lyase type I OS=*Synechococcus elongatus* (strain PCC 7942) GN=Synpcc7942_0112 PE=3 SV=1 |
| Q31KD6 | 1.915653 | 0.008992 | 0.00026 | 0.000498 | Type IV pilus assembly protein PilM OS=*Synechococcus elongatus* (strain PCC 7942) GN=Synpcc7942_2453 PE=4 SV=1 |
| Q31LW4 | 1.860608 | 0.004468 | 0.000127 | 0.000236 | Probable peptidase OS=*Synechococcus elongatus* (strain PCC 7942) GN=Synpcc7942_1925 PE=4 SV=1 |
| Q93AK0 | 1.846177 | 0.019625 | 8.61E-05 | 0.000159 | Cell division protein Ftn2 OS=*Synechococcus elongatus* (strain PCC 7942) GN=Synpcc7942_1943 PE=4 SV=1 |
| Q79PF2 | 1.83079 | 0.027555 | 0.000246 | 0.00045 | Glutathione peroxidase OS=*Synechococcus elongatus* (strain PCC 7942) GN=SEF0013 PE=3 SV=1 |
| Q31Q50 | 1.75903 | 0.004694 | 0.000298 | 0.000525 | Putative purple acid phosphatase OS=*Synechococcus elongatus* (strain PCC 7942) GN=Synpcc7942_0787 PE=4 SV=1 |
| Q31Q67 | 1.756009 | 0.001636 | 0.000183 | 0.000322 | Uncharacterized protein OS=*Synechococcus elongatus* (strain PCC 7942) GN=Synpcc7942_0770 PE=4 SV=1 |
| Q31S36 | 1.673385 | 0.017483 | 0.000184 | 0.000308 | Type 2 NADH dehydrogenase OS=*Synechococcus elongatus* (strain PCC 7942) GN=Synpcc7942_0101 PE=4 SV=1 |
| Q31QQ0 | 1.60873 | 0.013153 | 9.92E-05 | 0.00016 | Valine--tRNA ligase OS=*Synechococcus elongatus* (strain PCC 7942) GN=valS PE=3 SV=1 |
| Q99QJ5 | 1.582344 | 0.016321 | 0.00012 | 0.000189 | A LysR family protein OS=*Synechococcus elongatus* (strain PCC 7942) GN=rbcR PE=4 SV=1 |
| Q31LH0 | 1.540103 | 0.005251 | 0.000168 | 0.000259 | Fimbrial assembly protein PilC-like OS=*Synechococcus elongatus* (strain PCC 7942) GN=Synpcc7942_2069 PE=3 SV=1 |
| Q31RX6 | 1.489224 | 0.003584 | 0.000333 | 0.000496 | Chromophore lyase CpcS/CpeS OS=*Synechococcus elongatus* (strain PCC 7942) GN=cpcS PE=3 SV=1 |
| Q31RS1 | 1.488216 | 0.007536 | 8.87E-05 | 0.000132 | Putative zinc-binding oxidoreductase OS=*Synechococcus elongatus* (strain PCC 7942) GN=Synpcc7942_0216 PE=4 SV=1 |
| Q31QP2 | 1.436633 | 0.001432 | 9.67E-05 | 0.000139 | Uncharacterized protein OS=*Synechococcus elongatus* (strain |

| | | | | | |
|---|---|---|---|---|---|
| | | | | | PCC 7942) GN=Synpcc7942_0595 PE=4 SV=1 |
| Q31RM5 | -1.46069 | 0.006563 | 0.000351 | 0.00024 | Diaminopimelate decarboxylase OS=*Synechococcus elongatus* (strain PCC 7942) GN=lysA PE=3 SV=1 |
| Q8GJM5 | -1.56844 | 0.003017 | 0.000191 | 0.000122 | Putative uncharacterized protein SEN0014 OS=*Synechococcus elongatus* (strain PCC 7942) GN=SEN0014 PE=4 SV=1 |
| Q31KE3 | -1.61606 | 0.007863 | 9.83E-05 | 6.09E-05 | Methionine aminopeptidase OS=*Synechococcus elongatus* (strain PCC 7942) GN=map PE=3 SV=1 |
| Q31RF8 | -1.71068 | 0.024903 | 0.00012 | 7.02E-05 | Uncharacterized protein OS=*Synechococcus elongatus* (strain PCC 7942) GN=Synpcc7942_0329 PE=4 SV=1 |
| Q31JY4 | -1.72632 | 0.043695 | 0.000167 | 9.65E-05 | Uncharacterized protein OS=*Synechococcus elongatus* (strain PCC 7942) GN=Synpcc7942_2605 PE=4 SV=1 |
| Q31N14 | -1.79507 | 0.030187 | 0.000267 | 0.000149 | GTP-binding protein TypA OS=*Synechococcus elongatus* (strain PCC 7942) GN=Synpcc7942_1525 PE=4 SV=1 |
| Q31S56 | -1.8226 | 0.034798 | 0.000445 | 0.000244 | RNA-binding S4 OS=*Synechococcus elongatus* (strain PCC 7942) GN=Synpcc7942_0081 PE=4 SV=1 |
| Q31LQ3 | -1.90142 | 0.043926 | 0.000144 | 7.57E-05 | Processing protease OS=*Synechococcus elongatus* (strain PCC 7942) GN=Synpcc7942_1986 PE=3 SV=1 |
| Q31Q78 | -1.93134 | 0.032953 | 0.000105 | 5.46E-05 | Uncharacterized protein OS=*Synechococcus elongatus* (strain PCC 7942) GN=Synpcc7942_0759 PE=4 SV=1 |
| Q31M28 | -2.13602 | 0.023915 | 9.94E-05 | 4.65E-05 | Periplasmic binding protein of ABC transporter for natural amino acids OS=*Synechococcus elongatus* (strain PCC 7942) GN=Synpcc7942_1861 PE=4 SV=1 |
| Q31KF4 | -2.14307 | 0.019791 | 0.000174 | 8.13E-05 | Uncharacterized protein OS=*Synechococcus elongatus* (strain PCC 7942) GN=Synpcc7942_2435 PE=4 SV=1 |
| Q31MM4 | -2.23873 | 0.021095 | 0.000118 | 5.28E-05 | Probable oxidoreductase OS=*Synechococcus elongatus* (strain PCC 7942) GN=Synpcc7942_1665 PE=4 SV=1 |
| Q31ME3 | -2.25593 | 0.013491 | 0.000197 | 8.74E-05 | RNA polymerase sigma factor SigA2 OS=*Synechococcus elongatus* (strain PCC 7942) GN=sigA2 PE=1 SV=1 |
| Q31LA2 | -2.26231 | 0.030951 | 0.000121 | 5.35E-05 | Hydrogenobyrinic acid a,c-diamide cobaltochelatase OS=*Synechococcus elongatus* (strain PCC 7942) GN=Synpcc7942_2137 PE=4 SV=1 |

| | | | | | |
|---|---|---|---|---|---|
| Q935Y2 | -2.28522 | 0.0312 | 0.000113 | 4.92E-05 | Ribosomal protein S12 methylthiotransferase RimO OS=*Synechococcus elongatus* (strain PCC 7942) GN=rimO PE=3 SV=1 |
| Q31L97 | -2.36499 | 0.012003 | 0.000448 | 0.000189 | Translation initiation factor 1 (EIF-1/SUI1) OS=*Synechococcus elongatus* (strain PCC 7942) GN=Synpcc7942_2142 PE=4 SV=1 |
| Q31PF9 | -2.49891 | 0.010009 | 0.000205 | 8.21E-05 | Histidinol-phosphate aminotransferase OS=*Synechococcus elongatus* (strain PCC 7942) GN=hisC PE=3 SV=1 |

**Group 3 (Dark-adapted samples):**

Identifications satisfied both the automatic fold and statistical criteria in the Patternlab analysis.

| UniProt ID | Fold Change | p Value | Δ*glgC* | WT | Description |
|---|---|---|---|---|---|
| Q31RP3 | 4.187842 | 0.012435 | 0.000222 | 0.00093 | Alpha-1,4 glucan phosphorylase OS=*Synechococcus elongatus* (strain PCC 7942) GN=Synpcc7942_0244 PE=3 SV=1 |
| Q31KJ2 | 1.932827 | 0.013195 | 0.000362 | 0.000699 | Uncharacterized protein OS=*Synechococcus elongatus* (strain PCC 7942) GN=Synpcc7942_2397 PE=4 SV=1 |
| Q31S28 | 1.690714 | 0.022939 | 0.000482 | 0.000815 | DNA-binding ferritin-like protein (Oxidative damage protectant)-like OS=*Synechococcus elongatus* (strain PCC 7942) GN=Synpcc7942_0109 PE=3 SV=1 |
| Q54761 | 1.679318 | 0.042374 | 0.000351 | 0.00059 | Biotin carboxyl carrier protein OS=*Synechococcus elongatus* (strain PCC 7942) GN=accB PE=4 SV=1 |
| Q31S91 | 1.48368 | 0.024287 | 0.000434 | 0.000644 | Methylase involved in ubiquinone/menaquinone biosynthesis-like OS=*Synechococcus elongatus* (strain PCC 7942) GN=Synpcc7942_0046 PE=4 SV=1 |
| Q31KE0 | 1.449018 | 0.038153 | 0.001984 | 0.002875 | 1-Cys peroxiredoxin OS=*Synechococcus elongatus* (strain PCC 7942) GN=Synpcc7942_2449 PE=4 SV=1 |
| Q31RG9 | 1.436734 | 0.002948 | 0.000405 | 0.000582 | Uncharacterized protein OS=*Synechococcus elongatus* (strain PCC 7942) GN=Synpcc7942_0318 PE=4 SV=1 |
| Q31LJ0 | 1.434745 | 0.023697 | 0.000685 | 0.000982 | Photosystem I P700 chlorophyll a apoprotein A1 OS=*Synechococcus elongatus* (strain PCC 7942) GN=psaA PE=3 SV=1 |
| Q31KU2 | 1.409504 | 0.018316 | 0.002366 | 0.003334 | Transaldolase OS=*Synechococcus elongatus* (strain PCC 7942) GN=tal PE=3 SV=1 |
| P21577 | 1.353751 | 0.00217 | 0.00166 | 0.002248 | 6-phosphogluconate dehydrogenase, decarboxylating OS=*Synechococcus* |

| | | | | | |
|---|---|---|---|---|---|
| | | | | | *elongatus* (strain PCC 7942) GN=gnd PE=1 SV=4 |
| P29820 | 1.258397 | 0.00752 | 0.004173 | 0.005252 | Peptidyl-prolyl cis-trans isomerase OS=*Synechococcus elongatus* (strain PCC 7942) GN=rot PE=3 SV=1 |
| Q55041 | -1.30386 | 0.003691 | 0.000657 | 0.000504 | Phosphoribosylformylglycinamidine synthase subunit PurL OS=*Synechococcus elongatus* (strain PCC 7942) GN=purL PE=3 SV=3 |
| Q31QY6 | -1.31339 | 0.000798 | 0.000676 | 0.000515 | Nucleoside-diphosphate-sugar epimerases-like OS=*Synechococcus elongatus* (strain PCC 7942) GN=Synpcc7942_0501 PE=4 SV=1 |
| Q31L33 | -1.39192 | 0.006198 | 0.001269 | 0.000912 | 50S ribosomal protein L13 OS=*Synechococcus elongatus* (strain PCC 7942) GN=rplM PE=3 SV=1 |
| Q31QY1 | -1.39678 | 0.027284 | 0.000661 | 0.000473 | Uridylate kinase OS=*Synechococcus elongatus* (strain PCC 7942) GN=pyrH PE=3 SV=2 |
| Q31QJ2 | -1.39787 | 0.018374 | 0.00126 | 0.000902 | Glutamate-1-semialdehyde 2,1-aminomutase OS=*Synechococcus elongatus* (strain PCC 7942) GN=hemL PE=1 SV=3 |
| Q31L20 | -1.41685 | 0.017552 | 0.002574 | 0.001816 | 30S ribosomal protein S8 OS=*Synechococcus elongatus* (strain PCC 7942) GN=rpsH PE=3 SV=1 |
| Q31PV3 | -1.47048 | 0.006757 | 0.002324 | 0.00158 | 30S ribosomal protein S7 OS=*Synechococcus elongatus* (strain PCC 7942) GN=rpsG PE=3 SV=1 |
| Q31LL9 | -1.47288 | 0.00948 | 0.000612 | 0.000415 | Translation initiation factor IF-2 OS=*Synechococcus elongatus* (strain PCC 7942) GN=infB PE=3 SV=1 |
| Q31NR1 | -1.47833 | 0.019041 | 0.002116 | 0.001432 | 50S ribosomal protein L35 OS=*Synechococcus elongatus* (strain PCC 7942) GN=rpmI PE=3 SV=1 |
| P43087 | -1.50422 | 0.000117 | 0.000662 | 0.00044 | Delta-aminolevulinic acid dehydratase OS=*Synechococcus elongatus* (strain PCC 7942) GN=hemB PE=3 SV=2 |
| Q57311 | -1.54758 | 0.004817 | 0.000769 | 0.000497 | ORF 3 OS=*Synechococcus elongatus* (strain PCC 7942) GN=Synpcc7942_1429 PE=4 SV=1 |
| P0A3F4 | -1.5901 | 0.018767 | 0.014273 | 0.008976 | Nitrogen regulatory protein P-II OS=*Synechococcus elongatus* (strain PCC 7942) GN=glnB PE=1 SV=1 |
| Q31PX7 | -1.60468 | 0.01914 | 0.00075 | 0.000467 | Uncharacterized protein OS=*Synechococcus elongatus* (strain PCC 7942) GN=Synpcc7942_0862 PE=4 SV=1 |
| Q31RX1 | -1.66583 | 0.01148 | 0.000862 | 0.000518 | Uncharacterized protein OS=*Synechococcus elongatus* (strain PCC 7942) GN=Synpcc7942_0166 PE=4 SV=1 |
| Q31QE7 | -1.69044 | 0.038514 | 0.000678 | 0.000401 | ATP-dependent Clp protease adapter protein ClpS OS=*Synechococcus elongatus* (strain PCC 7942) GN=clpS PE=3 SV=1 |

| | | | | | |
|---|---|---|---|---|---|
| Q31RH4 | -1.70471 | 0.009279 | 0.000586 | 0.000344 | Uncharacterized protein OS=*Synechococcus elongatus* (strain PCC 7942) GN=Synpcc7942_0313 PE=4 SV=1 |
| O33698 | -1.7112 | 0.00507 | 0.000866 | 0.000506 | 30S ribosomal protein S1 OS=*Synechococcus elongatus* (strain PCC 7942) GN=rpsA PE=3 SV=1 |
| Q31QK2 | -1.73417 | 0.042149 | 0.001005 | 0.00058 | Transcription termination/antitermination protein NusG OS=*Synechococcus elongatus* (strain PCC 7942) GN=nusG PE=3 SV=1 |
| Q31RR1 | -1.75874 | 0.046613 | 0.000952 | 0.000541 | Sec-independent protein translocase protein TatA OS=*Synechococcus elongatus* (strain PCC 7942) GN=tatA PE=3 SV=1 |
| Q31NQ0 | -1.78082 | 0.010826 | 0.000729 | 0.000409 | Putative modulator of DNA gyrase OS=*Synechococcus elongatus* (strain PCC 7942) GN=Synpcc7942_1289 PE=4 SV=1 |
| Q31QL4 | -1.83192 | 0.021802 | 0.001086 | 0.000593 | Thioredoxin reductase OS=*Synechococcus elongatus* (strain PCC 7942) GN=Synpcc7942_0623 PE=3 SV=1 |
| Q31QK1 | -1.83757 | 0.015346 | 0.000616 | 0.000335 | Protein translocase subunit SecE OS=*Synechococcus elongatus* (strain PCC 7942) GN=secE PE=3 SV=1 |
| Q54760 | -1.84147 | 0.024151 | 0.000751 | 0.000408 | Elongation factor P OS=*Synechococcus elongatus* (strain PCC 7942) GN=efp PE=3 SV=1 |
| Q31LR4 | -1.96651 | 0.043726 | 0.000819 | 0.000417 | Uncharacterized protein OS=*Synechococcus elongatus* (strain PCC 7942) GN=Synpcc7942_1975 PE=4 SV=1 |
| Q31LW1 | -2.06855 | 0.008949 | 0.000767 | 0.000371 | Uncharacterized protein OS=*Synechococcus elongatus* (strain PCC 7942) GN=Synpcc7942_1928 PE=4 SV=1 |
| Q31L14 | -2.12946 | 0.032226 | 0.002565 | 0.001205 | 50S ribosomal protein L16 OS=*Synechococcus elongatus* (strain PCC 7942) GN=rplP PE=3 SV=1 |
| Q31KI8 | -2.38355 | 0.028787 | 0.003963 | 0.001663 | Heat shock protein Hsp20 OS=*Synechococcus elongatus* (strain PCC 7942) GN=Synpcc7942_2401 PE=3 SV=1 |

**Group 4 (Dark-adapted samples):**

These identifications were filtered out by the L-stringency in the Patternlab analysis.

| UniProt ID | Fold Change | p Value | Δ*glgC* | WT | Description |
|---|---|---|---|---|---|
| Q31M00 | 3.676863764 | 0.002787428 | 7.23E-05 | 0.000266 | Uncharacterized protein OS=*Synechococcus elongatus* (strain PCC 7942) GN=Synpcc7942_1889 PE=3 SV=1 |

| | | | | | |
|---|---|---|---|---|---|
| Q935Y7 | 2.445610472 | 0.006344068 | 0.000137498 | 0.000336 | Glycogen synthase OS=*Synechococcus elongatus* (strain PCC 7942) GN=glgA PE=3 SV=3 |
| Q54767 | 2.264607987 | 0.025292241 | 0.000120857 | 0.000274 | Probable short-chain dehydrogenase OS=*Synechococcus elongatus* (strain PCC 7942) GN=Synpcc7942_1596 PE=3 SV=1 |
| Q31QQ6 | 1.986179678 | 0.000781823 | 0.000281438 | 0.000559 | Uncharacterized protein OS=*Synechococcus elongatus* (strain PCC 7942) GN=Synpcc7942_0581 PE=4 SV=1 |
| P16954 | 1.912167026 | 0.004678815 | 0.000149736 | 0.000286 | 1,4-alpha-glucan branching enzyme GlgB OS=*Synechococcus elongatus* (strain PCC 7942) GN=glgB PE=1 SV=2 |
| Q31S93 | 1.85811386 | 0.039749429 | 5.24E-05 | 9.74E-05 | Uncharacterized protein OS=*Synechococcus elongatus* (strain PCC 7942) GN=Synpcc7942_0044 PE=4 SV=1 |
| Q31PW6 | 1.705913328 | 0.044262727 | 5.61E-05 | 9.57E-05 | Uncharacterized protein OS=*Synechococcus elongatus* (strain PCC 7942) GN=Synpcc7942_0873 PE=4 SV=1 |
| Q31R87 | 1.454908189 | 0.04416715 | 0.000191365 | 0.000278 | Anthranilate synthase, component II OS=*Synechococcus elongatus* (strain PCC 7942) GN=Synpcc7942_0400 PE=4 SV=1 |
| Q31PN8 | 1.42607167 | 0.015679039 | 0.000361614 | 0.000516 | Nicotinate-nucleotide pyrophosphorylase (Carboxylating) OS=*Synechococcus elongatus* (strain PCC 7942) GN=Synpcc7942_0951 PE=3 SV=1 |
| Q31RW4 | 1.413170147 | 0.011243906 | 6.25E-05 | 8.84E-05 | 3'-Phosphoadenosine 5'-phosphosulfate (PAPS) 3'-phosphatase-like OS=*Synechococcus elongatus* (strain PCC 7942) GN=Synpcc7942_0173 PE=4 SV=1 |
| Q31S89 | 1.392531756 | 0.009044063 | 4.06E-04 | 0.000566 | Pilin polypeptide PilA-like OS=*Synechococcus elongatus* (strain PCC 7942) GN=Synpcc7942_0048 PE=4 SV=1 |
| Q31LK6 | 1.383659764 | 0.025079631 | 0.000165188 | 0.000229 | Uncharacterized protein OS=*Synechococcus elongatus* (strain PCC 7942) GN=Synpcc7942_2033 PE=4 SV=1 |
| Q31S36 | -1.328518721 | 0.012852124 | 2.36E-04 | 0.000178 | Type 2 NADH dehydrogenase OS=*Synechococcus elongatus* (strain PCC 7942) GN=Synpcc7942_0101 PE=4 SV=1 |
| Q31N24 | -1.343260526 | 0.012127262 | 0.00056676 | 0.000422 | Protein serine/threonine phosphatase OS=*Synechococcus elongatus* (strain PCC 7942) GN=Synpcc7942_1515 PE=4 SV=1 |
| Q55267 | -1.374171263 | 0.013257134 | 0.000403758 | 0.000294 | ATP phosphoribosyltransferase regulatory subunit OS=*Synechococcus* |

| | | | | | |
|---|---|---|---|---|---|
| | | | | | *elongatus* (strain PCC 7942) GN=hisZ PE=3 SV=2 |
| Q31LV9 | -1.398184142 | 0.005711771 | 0.000102531 | 7.33E-05 | Uncharacterized protein OS=*Synechococcus elongatus* (strain PCC 7942) GN=Synpcc7942_1930 PE=4 SV=1 |
| Q31QC6 | -1.403868314 | 0.001037372 | 0.000464733 | 0.000331 | Carbamoyl-phosphate synthase (glutamine-hydrolyzing) OS=*Synechococcus elongatus* (strain PCC 7942) GN=Synpcc7942_0711 PE=3 SV=1 |
| Q8KPS9 | -1.413619626 | 0.016508977 | 0.000514673 | 0.000364 | Phycocyanobilin:ferredoxin oxidoreductase OS=*Synechococcus elongatus* (strain PCC 7942) GN=pcyA PE=3 SV=1 |
| Q31Q29 | -1.42883315 | 0.000455863 | 0.000354292 | 0.000248 | HAD-superfamily hydrolase subfamily IIB OS=*Synechococcus elongatus* (strain PCC 7942) GN=Synpcc7942_0808 PE=4 SV=1 |
| Q8GIS4 | -1.448384796 | 0.010585069 | 0.000447067 | 0.000309 | Photosystem II reaction center Psb28 protein OS=*Synechococcus elongatus* (strain PCC 7942) GN=psbW PE=3 SV=1 |
| Q8GIS1 | -1.460756084 | 0.042163766 | 0.00021763 | 0.000149 | Argininosuccinate lyase OS=*Synechococcus elongatus* (strain PCC 7942) GN=argH PE=3 SV=1 |
| Q31QS6 | -1.467336935 | 0.027127282 | 0.000298776 | 0.000204 | Uncharacterized protein OS=*Synechococcus elongatus* (strain PCC 7942) GN=Synpcc7942_0561 PE=4 SV=1 |
| Q31NP2 | -1.478943506 | 0.017672592 | 0.000457919 | 0.00031 | Malate dehydrogenase (Oxaloacetate decarboxylating) OS=*Synechococcus elongatus* (strain PCC 7942) GN=Synpcc7942_1297 PE=3 SV=1 |
| Q31PD2 | -1.479618939 | 0.027426425 | 0.000366407 | 0.000248 | Thiamine-phosphate synthase OS=*Synechococcus elongatus* (strain PCC 7942) GN=thiE PE=3 SV=1 |
| Q31KM0 | -1.487877292 | 0.026675987 | 9.39E-05 | 6.31E-05 | Hydrophobe/amphiphile efflux-1 HAE1 OS=*Synechococcus elongatus* (strain PCC 7942) GN=Synpcc7942_2369 PE=4 SV=1 |
| Q31QI3 | -1.49665918 | 0.005915582 | 1.40E-04 | 9.36E-05 | Photosystem I assembly protein Ycf4 OS=*Synechococcus elongatus* (strain PCC 7942) GN=ycf4 PE=3 SV=1 |
| Q31QR3 | -1.496819838 | 0.011400823 | 9.82E-05 | 6.56E-05 | Uncharacterized protein OS=*Synechococcus elongatus* (strain PCC 7942) GN=Synpcc7942_0574 PE=4 SV=1 |
| Q31KL8 | -1.498299047 | 0.001031121 | 9.98E-05 | 6.66E-05 | Acyl-[acyl-carrier-protein]--UDP-N-acetylglucosamine O-acyltransferase OS=*Synechococcus elongatus* (strain PCC 7942) GN=Synpcc7942_2371 PE=4 SV=1 |
| Q31QP9 | -1.516543429 | 0.006127401 | 0.000534159 | 0.000352 | Phosphoribosylglycinamide formyltransferase 2 |

| | | | | | |
|---|---|---|---|---|---|
| | | | | | OS=*Synechococcus elongatus* (strain PCC 7942) GN=purT PE=3 SV=1 |
| Q31PR2 | -1.51880447 | 0.046147831 | 0.00055492 | 0.000365 | Phosphoribosylaminoimidazole-succinocarboxamide synthase OS=*Synechococcus elongatus* (strain PCC 7942) GN=purC PE=3 SV=1 |
| Q31P62 | -1.551387792 | 0.0169002 | 0.000313742 | 0.000202 | Uncharacterized protein OS=*Synechococcus elongatus* (strain PCC 7942) GN=Synpcc7942_1127 PE=4 SV=1 |
| Q31RZ8 | -1.568401667 | 0.028639459 | 0.000369239 | 0.000235 | Acetolactate synthase OS=*Synechococcus elongatus* (strain PCC 7942) GN=Synpcc7942_0139 PE=3 SV=1 |
| Q31MA3 | -1.578498213 | 0.036500352 | 0.000184945 | 0.000117 | Uncharacterized protein OS=*Synechococcus elongatus* (strain PCC 7942) GN=Synpcc7942_1786 PE=4 SV=1 |
| Q31QV9 | -1.599273395 | 0.037332371 | 0.00019675 | 0.000123 | Lipopolysaccharide biosynthesis proteins LPS OS=*Synechococcus elongatus* (strain PCC 7942) GN=Synpcc7942_0528 PE=4 SV=1 |
| Q31NG7 | -1.623050142 | 0.03668784 | 0.000106888 | 6.59E-05 | Methionine synthase (B12-dependent) OS=*Synechococcus elongatus* (strain PCC 7942) GN=Synpcc7942_1372 PE=4 SV=1 |
| Q31PI4 | -1.626361048 | 0.044900777 | 0.000140949 | 8.67E-05 | HAD-superfamily hydrolase subfamily IA, variant 3 OS=*Synechococcus elongatus* (strain PCC 7942) GN=Synpcc7942_1005 PE=4 SV=1 |
| Q31NT0 | -1.632386329 | 0.007382131 | 3.42E-04 | 0.00021 | Arsenite-activated ATPase (ArsA) OS=*Synechococcus elongatus* (strain PCC 7942) GN=Synpcc7942_1259 PE=4 SV=1 |
| Q31MT0 | -1.638180818 | 0.020443858 | 5.12E-05 | 3.12E-05 | Protein splicing (Intein) site OS=*Synechococcus elongatus* (strain PCC 7942) GN=Synpcc7942_1609 PE=4 SV=1 |
| Q31PI2 | -1.641120335 | 0.022542099 | 0.000207138 | 0.000126 | Fumarate hydratase class II OS=*Synechococcus elongatus* (strain PCC 7942) GN=fumC PE=3 SV=1 |
| Q31M50 | -1.673808126 | 0.043869863 | 0.000119749 | 7.15E-05 | Uncharacterized protein OS=*Synechococcus elongatus* (strain PCC 7942) GN=Synpcc7942_1839 PE=4 SV=1 |
| Q31N14 | -1.678027707 | 0.013109246 | 3.07E-04 | 0.000183 | GTP-binding protein TypA OS=*Synechococcus elongatus* (strain PCC 7942) GN=Synpcc7942_1525 PE=4 SV=1 |
| Q31PP6 | -1.696016058 | 0.047334095 | 0.000327479 | 0.000193 | Acetylornithine aminotransferase OS=*Synechococcus elongatus* (strain PCC 7942) GN=argD PE=3 SV=1 |
| Q935Z2 | -1.699231939 | 0.026337301 | 0.00011312 | 6.66E-05 | Glycerol-3-phosphate dehydrogenase [NAD(P)+] OS=*Synechococcus* |

| | | | | | |
|---|---|---|---|---|---|
| | | | | | *elongatus* (strain PCC 7942) GN=gpsA PE=3 SV=3 |
| Q31M43 | -1.736201715 | 0.031445276 | 0.000374702 | 0.000216 | Uncharacterized protein OS=*Synechococcus elongatus* (strain PCC 7942) GN=Synpcc7942_1846 PE=4 SV=1 |
| Q31SC4 | -1.738633617 | 0.023300113 | 0.000471798 | 0.000271 | Uncharacterized protein OS=*Synechococcus elongatus* (strain PCC 7942) GN=Synpcc7942_0013 PE=4 SV=1 |
| Q31MF4 | -1.784800039 | 0.00300071 | 1.09E-04 | 6.12E-05 | Iron-regulated ABC transporter membrane component SufB OS=*Synechococcus elongatus* (strain PCC 7942) GN=Synpcc7942_1735 PE=4 SV=1 |
| Q31LY4 | -1.80479525 | 0.042418293 | 5.81E-05 | 3.22E-05 | Cyclic nucleotide-binding domain (CNMP-BD) protein OS=*Synechococcus elongatus* (strain PCC 7942) GN=Synpcc7942_1905 PE=4 SV=1 |
| Q31PH9 | -1.848620209 | 0.023618211 | 0.000442811 | 0.00024 | Ribosomal large subunit pseudouridine synthase B OS=*Synechococcus elongatus* (strain PCC 7942) GN=Synpcc7942_1010 PE=4 SV=1 |
| Q31K44 | -1.850624852 | 0.02708929 | 0.000349918 | 0.000189 | L-aspartate aminotransferase OS=*Synechococcus elongatus* (strain PCC 7942) GN=Synpcc7942_2545 PE=3 SV=1 |
| Q31QZ8 | -1.858588409 | 0.031396106 | 9.42E-05 | 5.07E-05 | Aldehyde dehydrogenase OS=*Synechococcus elongatus* (strain PCC 7942) GN=Synpcc7942_0489 PE=3 SV=1 |
| Q31K50 | -1.864245522 | 0.001666762 | 0.000186558 | 0.0001 | Uncharacterized protein OS=*Synechococcus elongatus* (strain PCC 7942) GN=Synpcc7942_2539 PE=4 SV=1 |
| Q31PF9 | -1.879210571 | 0.021117507 | 0.000163275 | 8.69E-05 | Histidinol-phosphate aminotransferase OS=*Synechococcus elongatus* (strain PCC 7942) GN=hisC PE=3 SV=1 |
| Q31KY6 | -1.881406581 | 0.015522385 | 0.000197576 | 0.000105 | Glutamate--cysteine ligase, putative OS=*Synechococcus elongatus* (strain PCC 7942) GN=Synpcc7942_2253 PE=4 SV=1 |
| Q31PN3 | -1.881551883 | 0.01374604 | 0.000474526 | 0.000252 | Uncharacterized protein OS=*Synechococcus elongatus* (strain PCC 7942) GN=Synpcc7942_0956 PE=4 SV=1 |
| P38045 | -1.883607495 | 0.032248169 | 0.000224979 | 0.000119 | Nitrate transport ATP-binding protein NrtC OS=*Synechococcus elongatus* (strain PCC 7942) GN=nrtC PE=3 SV=1 |
| Q31MD7 | -1.895271099 | 0.037860218 | 0.000168835 | 8.91E-05 | Armadillo:PBS lyase HEAT-like repeat OS=*Synechococcus elongatus* (strain PCC 7942) GN=Synpcc7942_1752 PE=4 SV=1 |

| | | | | | |
|---|---|---|---|---|---|
| Q31L36 | -1.912768225 | 0.043824266 | 0.000290119 | 0.000152 | Peptide chain release factor 1 OS=*Synechococcus elongatus* (strain PCC 7942) GN=prfA PE=3 SV=1 |
| Q31LB7 | -1.913388654 | 0.019216292 | 0.000172898 | 9.04E-05 | Carbamoyl-phosphate synthase small chain OS=*Synechococcus elongatus* (strain PCC 7942) GN=carA PE=3 SV=1 |
| Q31KQ9 | -1.927664655 | 0.032172295 | 0.000278209 | 0.000144 | C-terminal processing peptidase-2. Serine peptidase. MEROPS family S41A OS=*Synechococcus elongatus* (strain PCC 7942) GN=Synpcc7942_2330 PE=3 SV=1 |
| Q31L96 | -1.930790184 | 0.004024474 | 0.000311408 | 0.000161 | Tryptophan synthase beta chain OS=*Synechococcus elongatus* (strain PCC 7942) GN=trpB PE=3 SV=1 |
| Q31RB2 | -1.969922831 | 0.00626484 | 0.000136341 | 6.92E-05 | Processing protease OS=*Synechococcus elongatus* (strain PCC 7942) GN=Synpcc7942_0375 PE=3 SV=1 |
| Q31MT1 | -1.999981857 | 0.03087011 | 0.000136575 | 6.83E-05 | Mannose-1-phosphate guanylyltransferase (GDP) OS=*Synechococcus elongatus* (strain PCC 7942) GN=Synpcc7942_1608 PE=3 SV=1 |
| Q31PQ7 | -2.067594315 | 0.032473222 | 0.000116585 | 5.64E-05 | Lipid-A-disaccharide synthase OS=*Synechococcus elongatus* (strain PCC 7942) GN=Synpcc7942_0932 PE=4 SV=1 |
| Q31LC8 | -2.080269465 | 0.010563531 | 1.63E-04 | 7.82E-05 | Xylose repressor OS=*Synechococcus elongatus* (strain PCC 7942) GN=Synpcc7942_2111 PE=4 SV=1 |
| Q31M41 | -2.109516299 | 0.004507958 | 0.000362474 | 0.000172 | Aspartate-semialdehyde dehydrogenase OS=*Synechococcus elongatus* (strain PCC 7942) GN=asd PE=3 SV=1 |
| Q8GAA4 | -2.135171174 | 0.024229424 | 0.000363232 | 0.00017 | Uncharacterized protein OS=*Synechococcus elongatus* (strain PCC 7942) GN=sek0026 PE=4 SV=1 |
| Q31NU3 | -2.156414886 | 0.005211274 | 1.16E-04 | 5.39E-05 | Carotene isomerase OS=*Synechococcus elongatus* (strain PCC 7942) GN=Synpcc7942_1246 PE=4 SV=1 |
| Q31QJ6 | -2.236032453 | 0.026402499 | 6.00E-05 | 2.68E-05 | Succinate dehydrogenase subunit A OS=*Synechococcus elongatus* (strain PCC 7942) GN=Synpcc7942_0641 PE=4 SV=1 |
| Q31M06 | -2.286911381 | 0.017488142 | 0.000196195 | 8.58E-05 | Uncharacterized protein OS=*Synechococcus elongatus* (strain PCC 7942) GN=Synpcc7942_1883 PE=4 SV=1 |
| Q31LV5 | -2.306249976 | 0.037195886 | 0.000115561 | 5.01E-05 | D-alanyl-D-alanine carboxypeptidase/D-alanyl-D-alanine-endopeptidase OS=*Synechococcus elongatus* (strain PCC 7942) GN=Synpcc7942_1934 PE=4 SV=1 |

| | | | | | |
|---|---|---|---|---|---|
| Q31PJ1 | -2.373896667 | 0.039756219 | 0.000195387 | 8.23E-05 | ATP-dependent zinc metalloprotease FtsH OS=*Synechococcus elongatus* (strain PCC 7942) GN=ftsH PE=3 SV=1 |
| Q31M98 | -2.410920225 | 0.01909502 | 0.000103126 | 4.28E-05 | Uncharacterized protein OS=*Synechococcus elongatus* (strain PCC 7942) GN=Synpcc7942_1791 PE=4 SV=1 |
| Q31QL2 | -2.487332685 | 0.048161684 | 0.000163702 | 6.58E-05 | Single-stranded nucleic acid binding R3H OS=*Synechococcus elongatus* (strain PCC 7942) GN=Synpcc7942_0625 PE=4 SV=1 |
| Q31PN2 | -2.497225914 | 0.01276398 | 5.53E-04 | 0.000221 | Cob(I)yrinic acid a,c-diamide adenosyltransferase OS=*Synechococcus elongatus* (strain PCC 7942) GN=Synpcc7942_0957 PE=4 SV=1 |
| Q31PZ1 | -2.725454809 | 0.045474415 | 0.000133422 | 4.90E-05 | Twitching motility protein OS=*Synechococcus elongatus* (strain PCC 7942) GN=Synpcc7942_0847 PE=4 SV=1 |

Table S2. Plasmids and strains used in this study.

| Plasmids | Genotype | Reference |
|---|---|---|
| pXWK3_glgC | Δ*glgC*::KmR | This study |
| pXWK3-gnd | Δ*gnd*::KmR | This study |
| **Strains** | **Genotype** | **Reference** |
| *Synechococcus elongatus* PCC 7942 | Wild Type | |
| GlgC mutant | Δ*glgC* | This study |
| Gnd mutant | Δ*gnd* | This study |

Table S3. Oligonucleotides used for assembly of deletion mutants.

| Target gene | Oligonucleotides | PCR product | Assembled product |
|---|---|---|---|
| SYNPCC7942_RS03095 (deletion) | TTCCCGGCCGCGGCCAGTCAGCGCC<br>AGCCTG<br>TGCAGGTCGATCTCTCGACCACCCG<br>GCCGCT | Upstream | pXWK3_glgC |
| | AGACACAACGTGGCATTCTACGAGG<br>CCAATCTGGCG<br>TATATGAGTAAACTTGGTCTGACAG<br>TCGCGCGAGGAATGCCCT | Downstream | |
| | CTGTCAGACCAAGTTTACTCATATA<br>TACTTTAG<br>TACGGTTATCCACAGAATCAG | Vector | |
| | CCGGGTGGTCGAGAGATCGACCTGC<br>AGGGGG<br>GCCTCGTAGAATGCCACGTTGTGTC<br>TCAAAATCTC | KanR | |
| | TCCCCTGATTCTGTGGATAACCGTA<br>CGTGGAGACCGAAACCTTGCGCTC<br>TGGCGCTGACTGGCCGCGGCCGGGA<br>AGCCGA | GmR | |
| SYNPCC7942_RS00195 (deletion) | TTCCCGGCCGCGGTCCTTGACGGTC<br>TATAACC<br>TGCAGGTCGATCTAAGACATCATGC<br>AGTTCAC | Upstream | pXWK3_gnd |
| | AGACACAACGTGGTTGATTGAAATC<br>ACGGCG<br>GCGCGAGGAATGCAAGCCAACTGG<br>CGATCGAG | Downstream | |
| | GCCAGTTGGCTTGCATTCCTCGCGC<br>GACTGTCAGAC<br>GACCGTCAAGGACCGCGGCCGGGA<br>AGCCGA | Vector | |
| | GCATGATGTCTTAGATCGACCTGCA<br>GGGGG<br>GATTTCAATCAACCACGTTGTGTCTC<br>AAAATCTC | KanR | |

File S1. Map and sequence of pXWK3_glgC.

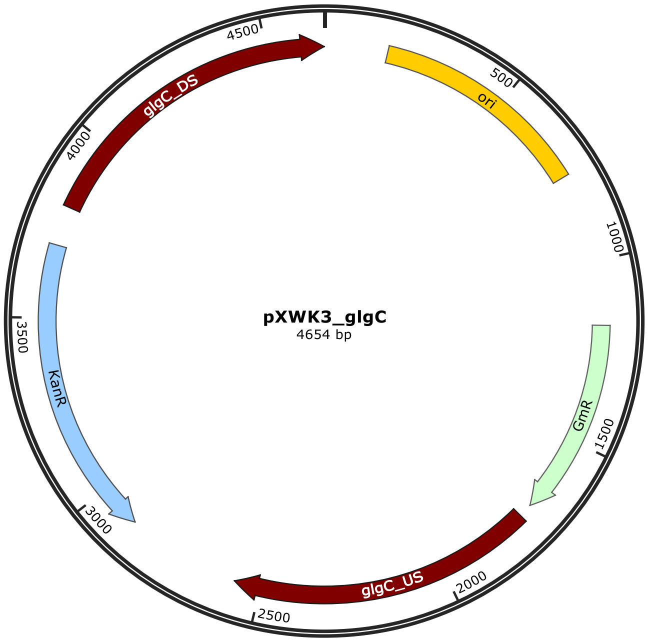


CTGTCAGACCAAGTTTACTCATATATACTTTAGATTGATTTAAAACTTCATTTTTAATTTAAAAGGATCT
AGGTGAAGATCCTTTTTGATAATCTCATGACCAAAATCCCTTAACGTGAGTTTTCGTTCCACTGAGCGTC
AGACCCCGTAGAAAAGATCAAAGGATCTTCTTGAGATCCTTTTTTTCTGCGCGTAATCTGCTGCTTGCAA
ACAAAAAAACCACCGCTACCAGCGGTGGTTTGTTTGCCGGATCAAGAGCTACCAACTCTTTTTCCGAAGG
TAACTGGCTTCAGCAGAGCGCAGATACCAAATACTGTTCTTCTAGTGTAGCCGTAGTTAGGCCACCACTT
CAAGAACTCTGTAGCACCGCCTACATACCTCGCTCTGCTAATCCTGTTACCAGTGGCTGCTGCCAGTGGC
GATAAGTCGTGTCTTACCGGGTTGGACTCAAGACGATAGTTACCGGATAAGGCGCAGCGGTCGGGCTGAA
CGGGGGGTTCGTGCACACAGCCCAGCTTGGAGCGAACGACCTACACCGAACTGAGATACCTACAGCGTGA
GCTATGAGAAAGCGCCACGCTTCCCGAAGGGAGAAAGGCGGACAGGTATCCGGTAAGCGGCAGGGTCGGA
ACAGGAGAGCGCACGAGGGAGCTTCCAGGGGGAAACGCCTGGTATCTTTATAGTCCTGTCGGGTTTCGCC
ACCTCTGACTTGAGCGTCGATTTTTGTGATGCTCGTCAGGGGGGCGGAGCCTATGGAAAAACGCCAGCAA
CGCGGCCTTTTTACGGTTCCTGGCCTTTTGCTGGCCTTTTGCTCACATGTTCTTTCCTGCGTTATCCCCT
GATTCTGTGGATAACCGTACGTGGAGACCGAAACCTTGCGCTCGTTCGCCAGCCAGGACAGAAATGCCTC
GACTTCGCTGCTGCCCAAGGTTGCCGGGTGACGCACACCGTGGAAACGGATGAAGGCACGAACCCAGTTG
ACATAAGCCTGTTCGGTTCGTAAACTGTAATGCAAGTAGCGTATGCGCTCACGCAACTGGTCCAGAACCT
TGACCGAACGCAGCGGTGGTAACGGCGCAGTGGCGGTTTTCATGGCTTGTTATGACTGTTTTTTTGTACA
GTCTATGCCTCGGGCATCCAAGCAGCAAGCGCGTTACGCCGTGGGTCGATGTTTGATGTTATGGAGCAGC
AACGATGTTACGCAGCAGGGCAGTCGCCCTAAAACAAAGTTAGGTGGCTCAAGTATGGGCATCATTCGCA
CATGTAGGCTCGGCCCTGACCAAGTCAAATCCATGCGGGCTGCTCTTGATCTTTTCGGTCGTGAGTTCGG
AGACGTAGCCACCTACTCCCAACATCAGCCGGACTCCGATTACCTCGGGAACTTGCTCCGTAGTAAGACA
TTCATCGCGCTTGCTGCCTTCGACCAAGAAGCGGTTGTTGGCGCTCTCGCGGCTTACGTTCTGCCCAAGT
TTGAGCAGCCGCGTAGTGAGATCTATATCTATGATCTCGCAGTCTCCGGCGAGCACCGGAGGCAGGGCAT
TGCCACCGCGCTCATCAATCTCCTCAAGCATGAGGCCAACGCGCTTGGTGCTTATGTGATCTACGTGCAA
GCAGATTACGGTGACGATCCCGCAGTGGCTCTCTATACAAAGTTGGGCATACGGAAGAAGTGATGCACT

TTGATATCGACCCAAGTACCGCCACCTAACAATTCGTTCAAGCCGAGATCGGCTTCCCGGCCGCGGCCAG
TCAGCGCCAGCCTGGTCAACAGCGCGGACATCGTCGCCCAAGCGGCTGAAATCCGCTGACAGAATGGACG
GAGCAATAACAACGGATTTTTGCGTCATAGCCTTTGGACTGAAGCGGCGATCGCTCCTCAGTTTACCAAC
AGCTGGTGCCCGCGCCATTGTCCGTAGCCTTGGTTACCTCCGATCGATCCCGGAGCTGCGTAGGGTAAGA
AATCGCAAAGGCACAGTTTGCTTGCCAAGACTGACTCACCGGCAGACACTGGGTCTCAGCGCCGCTTGAC
TTGGGGGAGATTGATTGTGAAAAACGTGCTGGCGATCATTCTCGGTGGAGGCGCAGGCAGTCGTCTCTAT
CCACTAACCAAACAGCGCGCCAAACCAGCGGTCCCCCTGGCGGGCAAATACCGCTTGATCGATATTCCCG
TCAGCAATTGCATCAACGCTGACATCAACAAAATCTATGTGCTGACGCAGTTTAACTCTGCCTCGCTCAA
CCGCCACCTCAGTCAGACCTACAACCTCTCCAGCGGCTTTGGCAATGGCTTTGTTGAGGTGCTAGCAGCT
CAGATTACGCCGGAGAACCCCAACTGGTTCCAAGGCACCGCCGATGCGGTTCGCCAGTATCTCTGGCTAA
TCAAAGAGTGGGATGTGGATGAGTACCTGATCCTGTCGGGGGATCATCTCTACCGCATGGACTATAGCCA
GTTCATTCAGCGGCACCGAGACACCAATGCCGACATCACACTCTCGGTCTTGCCGATCGATGAAAAGCGC
GCCTCTGATTTTGGCCTGATGAAGCTAGATGGCAGCGGCCGGGTGGTCGAGAGATCGACCTGCAGGGGGG
GGGGGGCGCTGAGGTCTGCCTCGTGAAGAAGGTGTTGCTGACTCATACCAGGCCTGAATCGCCCCATCAT
CCAGCCAGAAAGTGAGGGAGCCACGGTTGATGAGAGCTTTGTTGTAGGTGGACCAGTTGGTGATTTTGAA
CTTTTGCTTTGCCACGGAACGGTCTGCGTTGTCGGGAAGATGCGTGATCTGATCCTTCAACTCAGCAAAA
GTTCGATTTATTCAACAAAGCCGCCGTCCCGTCAAGTCAGCGTAATGCTCTGCCAGTGTTACAACCAATT
AACCAATTCTGATTAGAAAAACTCATCGAGCATCAAATGAAACTGCAATTTATTCATATCAGGATTATCA
ATACCATATTTTTGAAAAAGCCGTTTCTGTAATGAAGGAGAAAACTCACCGAGGCAGTTCCATAGGATGG
CAAGATCCTGGTATCGGTCTGCGATTCCGACTCGTCCAACATCAATACAACCTATTAATTTCCCCTCGTC
AAAAATAAGGTTATCAAGTGAGAAATCACCATGAGTGACGACTGAATCCGGTGAGAATGGCAAAAGCTTA
TGCATTTCTTTCCAGACTTGTTCAACAGGCCAGCCATTACGCTCGTCATCAAAATCACTCGCATCAACCA
AACCGTTATTCATTCGTGATTGCGCCTGAGCGAGACGAAATACGCGATCGCTGTTAAAAGGACAATTACA
AACAGGAATCGAATGCAACCGGCGCAGGAACACTGCCAGCGCATCAACAATATTTTCACCTGAATCAGGA
TATTCTTCTAATACCTGGAATGCTGTTTTCCCGGGGATCGCAGTGGTGAGTAACCATGCATCATCAGGAG
TACGGATAAAATGCTTGATGGTCGGAAGAGGCATAAATTCCGTCAGCCAGTTTAGTCTGACCATCTCATC
TGTAACATCATTGGCAACGCTACCTTTGCCATGTTTCAGAAACAACTCTGGCGCATCGGGCTTCCCATAC
AATCGATAGATTGTCGCACCTGATTGCCCGACATTATCGCGAGCCCATTTATACCCATATAAATCAGCAT
CCATGTTGGAATTTAATCGCGGCCTCGAGCAAGACGTTTCCCGTTGAATATGGCTCATAACACCCCTTGT
ATTACTGTTTATGTAAGCAGACAGTTTTATTGTTCATGATGATATATTTTTATCTTGTGCAATGTAACAT
CAGAGATTTTGAGACACAACGTGGCATTCTACGAGGCCAATCTGGCGCTGACTCAGCAACCTAGCCCACC
CTTCAGCTTCTACGACGAGCAGGCGCCGATTTACACCCGCGCTCGCTACCTGCCGCCAACCAAGCTGCTC
GATTGCCAGGTGACCCAGTCGATCATTGGCGAGGGCTGCATTCTCAAGCAATGCACCGTTCAGAATTCCG
TCTTAGGGATTCGCTCCCGCATTGAGGCCGACTGCGTGATCCAGGACGCCTTGTTGATGGGCGCTGACTT
CTACGAAACCTCGGAGCTACGGCACCAGAATCGGGCCAATGGCAAAGTGCCGATGGGAATCGGCAGTGGC
AGCACCATCCGTCGCGCCATCGTCGACAAAAATGCCCACATTGGCCAGAACGTTCAGATCGTCAACAAAG
ACCATGTGGAAGAGGCCGATCGCGAAGATCTGGGCTTTATGATCCGCAGCGGCATTGTCGTTGTGGTCAA
AGGGGCGGTTATTCCCGACAACACGGTGATCTAACACCATGCGCCTCGGCAAAGTTGTCAAATCCAACTC
GCACTGTGACTACGTTGTCCAAGTGGATGACGACATGGATGTCTCCAATCCACCAGCCGCCGAGAGCTAT
GGATTTGGCAGCTTTGTTCGTTTGGAAGGCGATCGCCATTGGGCGATCGGAGTGGTCTACAACTCGCAGT
TGTTTAATCCCCTCTTTCTGAACAACGGGCCGCGTCTTTCAAGTGGGCCCGACCCACTATTCACGCCTGA
TTTAATCAATGAAACCCGCACGCTGCTCTCCACGGTTTTGATCGGCAGCTTAGCGGAAACAGAGGGCGGC
GATCGCTACGGCGAACAGGGCATTCCTCGCGCGA

File S2. Map and sequence of pXWK3_gnd.

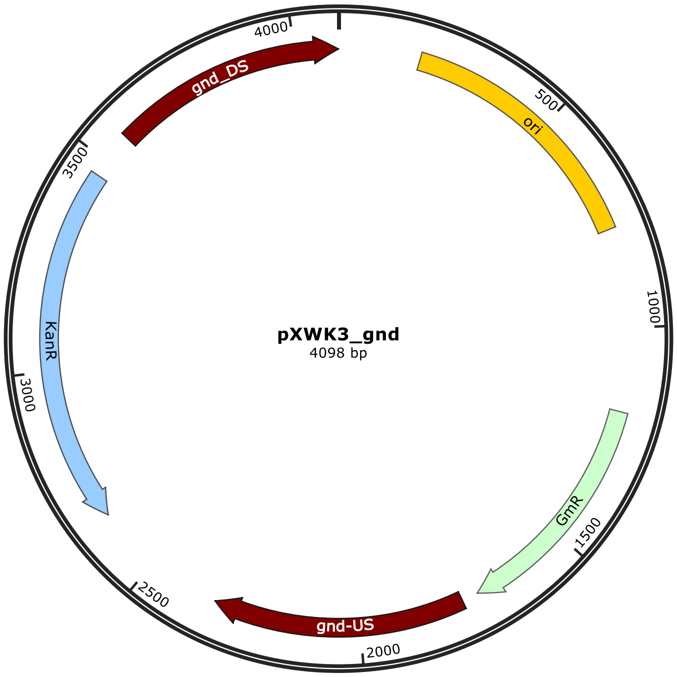


GCATTCCTCGCGCGACTGTCAGACCAAGTTTACTCATATATACTTTAGATTGATTTAAAACTTCATTTTT
AATTTAAAAGGATCTAGGTGAAGATCCTTTTTGATAATCTCATGACCAAAATCCCTTAACGTGAGTTTTC
GTTCCACTGAGCGTCAGACCCCGTAGAAAAGATCAAAGGATCTTCTTGAGATCCTTTTTTTCTGCGCGTA
ATCTGCTGCTTGCAAACAAAAAAACCACCGCTACCAGCGGTGGTTTGTTTGCCGGATCAAGAGCTACCAA
CTCTTTTTCCGAAGGTAACTGGCTTCAGCAGAGCGCAGATACCAAATACTGTTCTTCTAGTGTAGCCGTA
GTTAGGCCACCACTTCAAGAACTCTGTAGCACCGCCTACATACCTCGCTCTGCTAATCCTGTTACCAGTG
GCTGCTGCCAGTGGCGATAAGTCGTGTCTTACCGGGTTGGACTCAAGACGATAGTTACCGGATAAGGCGC
AGCGGTCGGGCTGAACGGGGGGTTCGTGCACACAGCCCAGCTTGGAGCGAACGACCTACACCGAACTGAG
ATACCTACAGCGTGAGCTATGAGAAAGCGCCACGCTTCCCGAAGGGAGAAAGGCGGACAGGTATCCGGTA
AGCGGCAGGGTCGGAACAGGAGAGCGCACGAGGGAGCTTCCAGGGGGAAACGCCTGGTATCTTTATAGTC
CTGTCGGGTTTCGCCACCTCTGACTTGAGCGTCGATTTTTGTGATGCTCGTCAGGGGGGCGGAGCCTATG
GAAAAACGCCAGCAACGCGGCCTTTTTACGGTTCCTGGCCTTTTGCTGGCCTTTTGCTCACATGTTCTTT
CCTGCGTTATCCCCTGATTCTGTGGATAACCGTACGTGGAGACCGAAACCTTGCGCTCGTTCGCCAGCCA
GGACAGAAATGCCTCGACTTCGCTGCTGCCCAAGGTTGCCGGGTGACGCACACCGTGGAAACGGATGAAG
GCACGAACCCAGTTGACATAAGCCTGTTCGGTTCGTAAACTGTAATGCAAGTAGCGTATGCGCTCACGCA
ACTGGTCCAGAACCTTGACCGAACGCAGCGGTGGTAACGGCGCAGTGGCGGTTTTCATGGCTTGTTATGA
CTGTTTTTTTGTACAGTCTATGCCTCGGGCATCCAAGCAGCAAGCGCGTTACGCCGTGGGTCGATGTTTG
ATGTTATGGAGCAGCAACGATGTTACGCAGCAGGGCAGTCGCCCTAAAACAAAGTTAGGTGGCTCAAGTA
TGGGCATCATTCGCACATGTAGGCTCGGCCCTGACCAAGTCAAATCCATGCGGGCTGCTCTTGATCTTTT
CGGTCGTGAGTTCGGAGACGTAGCCACCTACTCCCAACATCAGCCGGACTCCGATTACCTCGGGAACTTG
CTCCGTAGTAAGACATTCATCGCGCTTGCTGCCTTCGACCAAGAAGCGGTTGTTGGCGCTCTCGCGGCTT
ACGTTCTGCCCAAGTTTGAGCAGCCGCGTAGTGAGATCTATATCTATGATCTCGCAGTCTCCGGCGAGCA
CCGGAGGCAGGGCATTGCCACCGCGCTCATCAATCTCCTCAAGCATGAGGCCAACGCGCTTGGTGCTTAT
GTGATCTACGTGCAAGCAGATTACGGTGACGATCCCGCAGTGGCTCTCTATACAAAGTTGGGCATACGGG

AAGAAGTGATGCACTTTGATATCGACCCAAGTACCGCCACCTAACAATTCGTTCAAGCCGAGATCGGCTT
CCCGGCCGCGGTCCTTGACGGTCTATAACCGCACGGCCGAAAAGACTGAAGCGTTCATGGCCGATCGCGC
CCAAGGCAAGAACATTGTGCCGGCTTACAGTCTGGAAGACTTTGTTGCCAGCTTGGAACGTCCGCGCCGC
ATTTTGGTGATGGTTAAAGCGGGCGGCCCGGTGGATGCCGTGGTCGAGCAGCTCAAACCCCTGCTAGATC
CCGGTGACTTGATCATCGATGGGGGCAACTCACTGTTTACCGACACCGAGCGCCGCGTCAAAGATCTAGA
AGCGCTAGGTCTGGGTTTCATGGGCATGGGCGTCAGTGGTGGCGAAGAGGGGGCTTTGAATGGCCCCAGT
TTGATGCCGGGGGGCACCCAAGCGGCCTACGAAGCCGTGGAGCCGATCGTGCGCAGCATTGCTGCCCAAG
TCGACGATGGCCCCTGCGTTACCTACATCGGTCCTGGTGGCTCTGGGCACTACGTCAAGATGGTGCACAA
CGGCATTGAATATGGCGATATGCAGCTGATCGCCGAAGCCTATGACCTCCTCAAATCGGTGGCAGGCCTC
AATGCCAGTGAACTGCATGATGTCTTAGATCGACCTGCAGGGGGGGGGGGGCGCTGAGGTCTGCCTCGTG
AAGAAGGTGTTGCTGACTCATACCAGGCCTGAATCGCCCCATCATCCAGCCAGAAAGTGAGGGAGCCACG
GTTGATGAGAGCTTTGTTGTAGGTGGACCAGTTGGTGATTTTGAACTTTTGCTTTGCCACGGAACGGTCT
GCGTTGTCGGGAAGATGCGTGATCTGATCCTTCAACTCAGCAAAAGTTCGATTTATTCAACAAAGCCGCC
GTCCCGTCAAGTCAGCGTAATGCTCTGCCAGTGTTACAACCAATTAACCAATTCTGATTAGAAAAACTCA
TCGAGCATCAAATGAAACTGCAATTTATTCATATCAGGATTATCAATACCATATTTTTGAAAAAGCCGTT
TCTGTAATGAAGGAGAAAACTCACCGAGGCAGTTCCATAGGATGGCAAGATCCTGGTATCGGTCTGCGAT
TCCGACTCGTCCAACATCAATACAACCTATTAATTTCCCCTCGTCAAAAATAAGGTTATCAAGTGAGAAA
TCACCATGAGTGACGACTGAATCCGGTGAGAATGGCAAAAGCTTATGCATTTCTTTCCAGACTTGTTCAA
CAGGCCAGCCATTACGCTCGTCATCAAAATCACTCGCATCAACCAAACCGTTATTCATTCGTGATTGCGC
CTGAGCGAGACGAAATACGCGATCGCTGTTAAAAGGACAATTACAAACAGGAATCGAATGCAACCGGCGC
AGGAACACTGCCAGCGCATCAACAATATTTTCACCTGAATCAGGATATTCTTCTAATACCTGGAATGCTG
TTTTCCCGGGGATCGCAGTGGTGAGTAACCATGCATCATCAGGAGTACGGATAAAATGCTTGATGGTCGG
AAGAGGCATAAATTCCGTCAGCCAGTTTAGTCTGACCATCTCATCTGTAACATCATTGGCAACGCTACCT
TTGCCATGTTTCAGAAACAACTCTGGCGCATCGGGCTTCCCATACAATCGATAGATTGTCGCACCTGATT
GCCCGACATTATCGCGAGCCCATTTATACCCATATAAATCAGCATCCATGTTGGAATTTAATCGCGGCCT
CGAGCAAGACGTTTCCCGTTGAATATGGCTCATAACACCCCTTGTATTACTGTTTATGTAAGCAGACAGT
TTTATTGTTCATGATGATATATTTTTATCTTGTGCAATGTAACATCAGAGATTTTGAGACACAACGTGGT
TGATTGAAATCACGGCGGATATTTTCACCAAAGTCGATGACTTGGGTACTGGTCAGCCCTTGGTCGAGCT
GATTTTAGATGCCGCTGGTCAAAAAGGAACCGGTCGCTGGACGGTGGAAACGGCACTGGAAATTGGCGTT
GCGATTCCAACCATCATTGCTGCTGTCAACGCCCGCATCCTGTCCTCGATCAAAGCCGAGCGTCAGGCAG
CTTCTGAAATTCTTTCAGGACCGATTACCGAGCCCTTCAGTGGCGATCGCCAAGCCTTTATCGACAGTGT
GCGCGATGCGCTCTACTGCTCGAAAATTTGCTCCTATGCCCAAGGCATGGCGCTGCTAGCCAAAGCCTCC
CAGGTCTACAACTACGGTCTGAATTTAGGTGAACTGGCGCGGATTTGGAAAGGCGGCTGCATCATTCGGG
CGGGTTTCCTCAACAAGATTAAGCAGGCCTATGATGCTGACCCAACGCTGGCGAATCTGTTGCTGGCACC
CGAATTCCGGCAGACGATTCTCGATCGCCAGTTGGCTT
